# Taxonomy, phylogeny and biogeography of African spurfowls (Galliformes, Phasianidae, Coturnicinae, *Pternistis* spp.)

**DOI:** 10.1101/329243

**Authors:** Tshifhiwa G. Mandiwana-Neudani, Robin M. Little, Timothy M. Crowe, Rauri C.K. Bowie

## Abstract

During much of the 20^th^ Century, partridge/quail-like, Afro-Asian phasianine birds referred to commonly as African spurfowls, francolins and/or partridges had a tortuous taxonomic history. Because of striking autapomorphic differences in plumage, vocalizations and ecology in some of these taxa, as many as nine genera and nearly 200 clinal and/or idiosyncratic subspecies, embedded within a polyphyletic Perdicinae, were recognized. In 1963, two clades, 28 ‘francolin’ and ‘spurfowl’ species (*fisante* in Afrikaans) and 13 Afro-Asiatic ‘francolins’ and ‘partridges’ (*patryse* in Afrikaans), were combined into a single genus (*Francolinus*) – the largest within the Galliformes – comprising about 100 subspecies. Furthermore, *fisante* and *patryse* were partitioned into several unnamed “Groups” and four “Unplaced” species. Here, we use morphological, behavioural and DNA evidence to produce: a comprehensive revision of the taxonomy and phylogeny of the *fisante* clade; a stable classification system for tis component taxa; and hypotheses vis-à-vis eco-biogeographical processes that promoted their speciation and cladogenesis. We shift *Francolinus* spp. sensu stricto (members of the Spotted Group) and the Unplaced ‘*Francolinus’ gularis* from the *fisante* clade to the *patryse* [discussed in Mandiwana-Neudani et al., in review]. An Unplaced *fisant, ‘F.’ nahani*, is linked with *Ptilopachus petrosus* (another African endemic ‘partridge‘) within the Odontophoridae (New World ‘Quails‘). We recognize 25 species of fisante (hereafter spurfowls), including seven with subspecies. They comprise 34 terminal taxa placed within a single genus, *Pternistis*, sister to *Ammoperdix* and *Perdicula* spp., *Coturnix* ‘quails’ and *Alectoris* ‘partridges‘, within the now monophyletic Coturnicinae. Only one of four putative Groups of spurfowls, the Bare-throated Group, is monophyletic. The other three Groups (Montane, Scaly and Vermiculated) are para- or polyphyletic. Several species pairs of spurfowls, most notably *P. afer* and *cranchii*, hybridize in para/sympatry. One Bare-throated spurfowl, *P. rufopictus*, may be the product of stabilized hybridization between *P. afer* and/or *cranchii* and *P. leucoscepus*.

## Introduction

During much of the 20^th^ Century, there was little consensus relating to the taxonomy and phylogeny of Afro-Asian quail and partridge-like galliforms within the Phasianidae, variously commonly known as francolins, spurfowls and/or partridges. As many as nine genera [1] and nearly 200 clinal and/or idiosyncratic subspecies [2], embedded within a polyphyletic Perdicinae [3] were recognized. In 1963, Mrs B.P. ‘Pat’ Hall comprehensively revised the taxonomy of many of these taxa [2]. She argued convincingly that they should be combined within a single ‘mega-genus‘, *Francolinus*, comprising 41 species - the largest genus in the order Galliformes and the twelfth largest in Aves [4]. Thirty-six of these species are endemic to Africa, five to Asia. Hall also synonymized many subspecies, reducing the nearly 200 to just over 100 [2].

However, literally immediately after this ‘lumping/ synonymizing’ exercise, Hall divided “francolins” into two, unnamed, putatively monophyletic major clades, comprised of eight (also unnamed and putatively monophyletic) “Groups” and four “Unplaced” species [2]. The major clades of francolins correspond to what Afrikaans-speakers commonly refer to as *fisante* (’pheasants‘) and *patryse* (’partridges‘) [5, 6]. We deal with the *patryse* elsewhere [Mandiwana-Neudani et al., in review]. Hall‘s *fisante* (hereafter spurfowls) included an Asiatic Spotted Group (incorporating the nominate species *F. francolinus* and congeners), four other Groups (Vermiculated, Montane, Scaly and Bare-throated) and two Unplaced species (*nahani* and *gularis*) [2]. Morphologically, these taxa generally have: plain or plain-vermiculated back-plumage; brown/black/red tarsi with long – sometimes multiple – spurs; emit raucous, grating vocalizations; and roost/perch in large bushes or trees [6]. Members of one Group of spurfowls (the Bare-throated), differ from the others in having bare, brightly coloured skin around the eye and/or on the throat [2].

Within the spurfowls, Hall [2] recognized 28 species, which are generally sexually monomorphic, with males (and females of some taxa) of most species having at least a single (often two), long tarsal spurs. The species differ markedly in plumage, ecology, behaviour and distributional patterns [7, 8, 9].

**Table 1.**
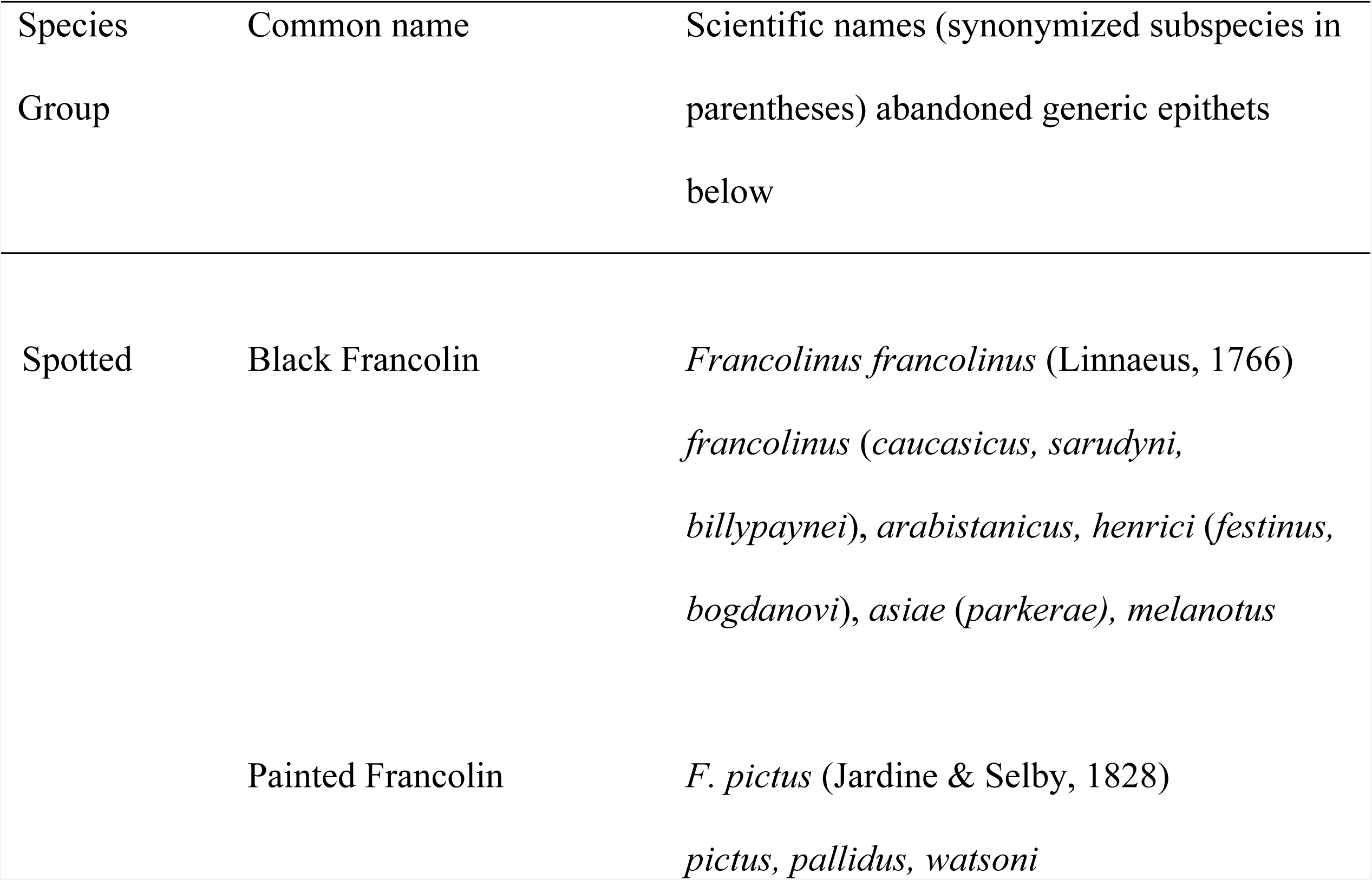
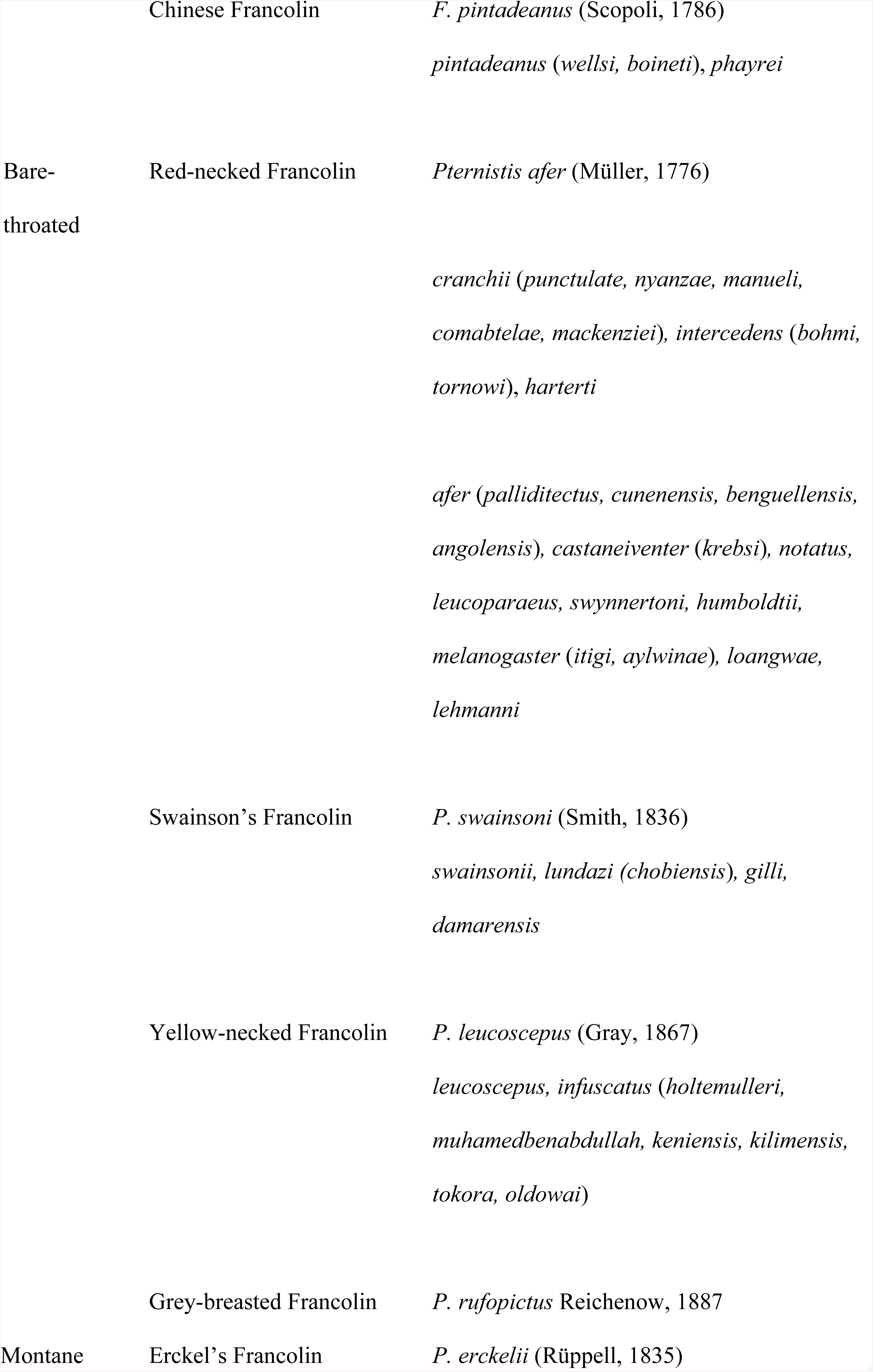
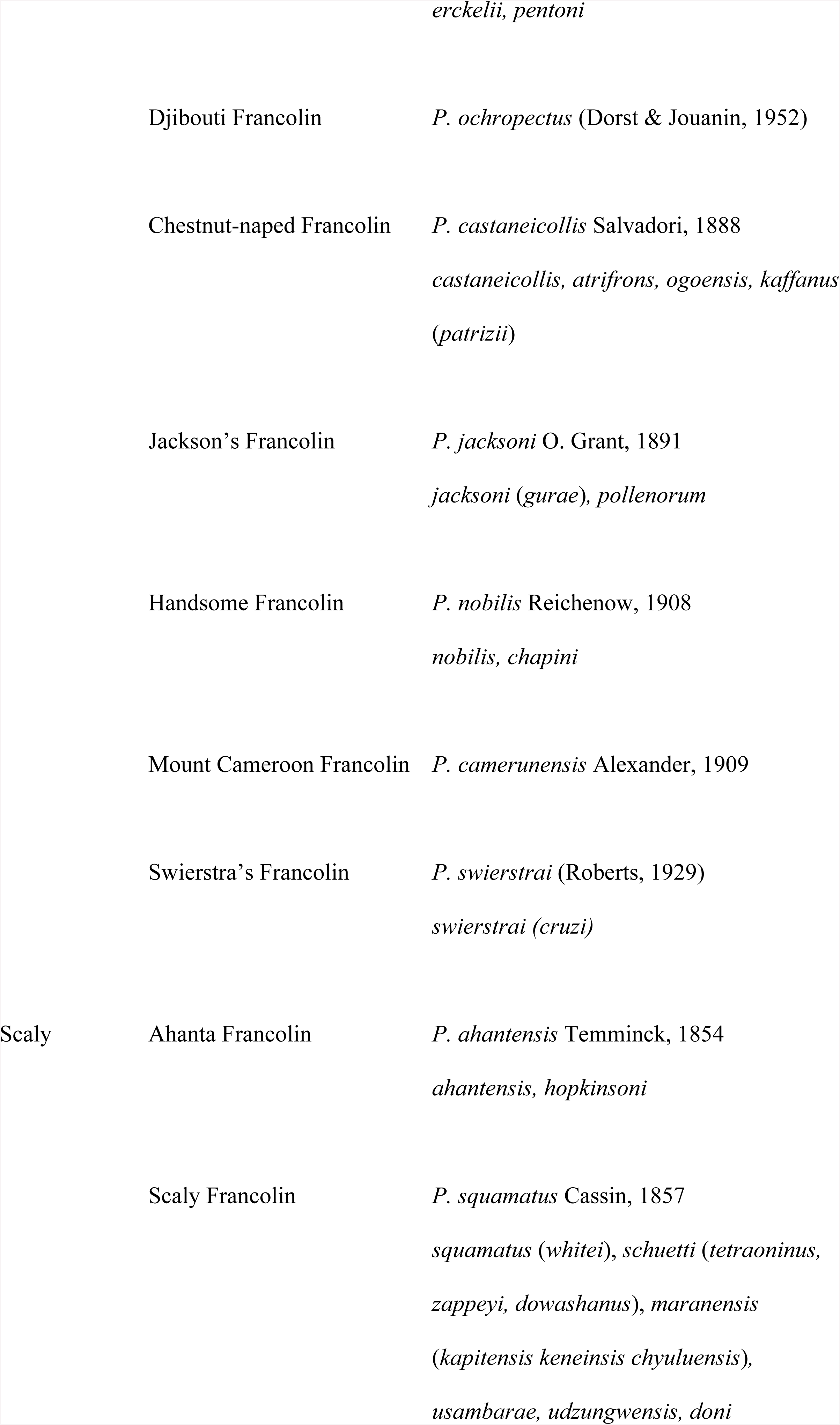
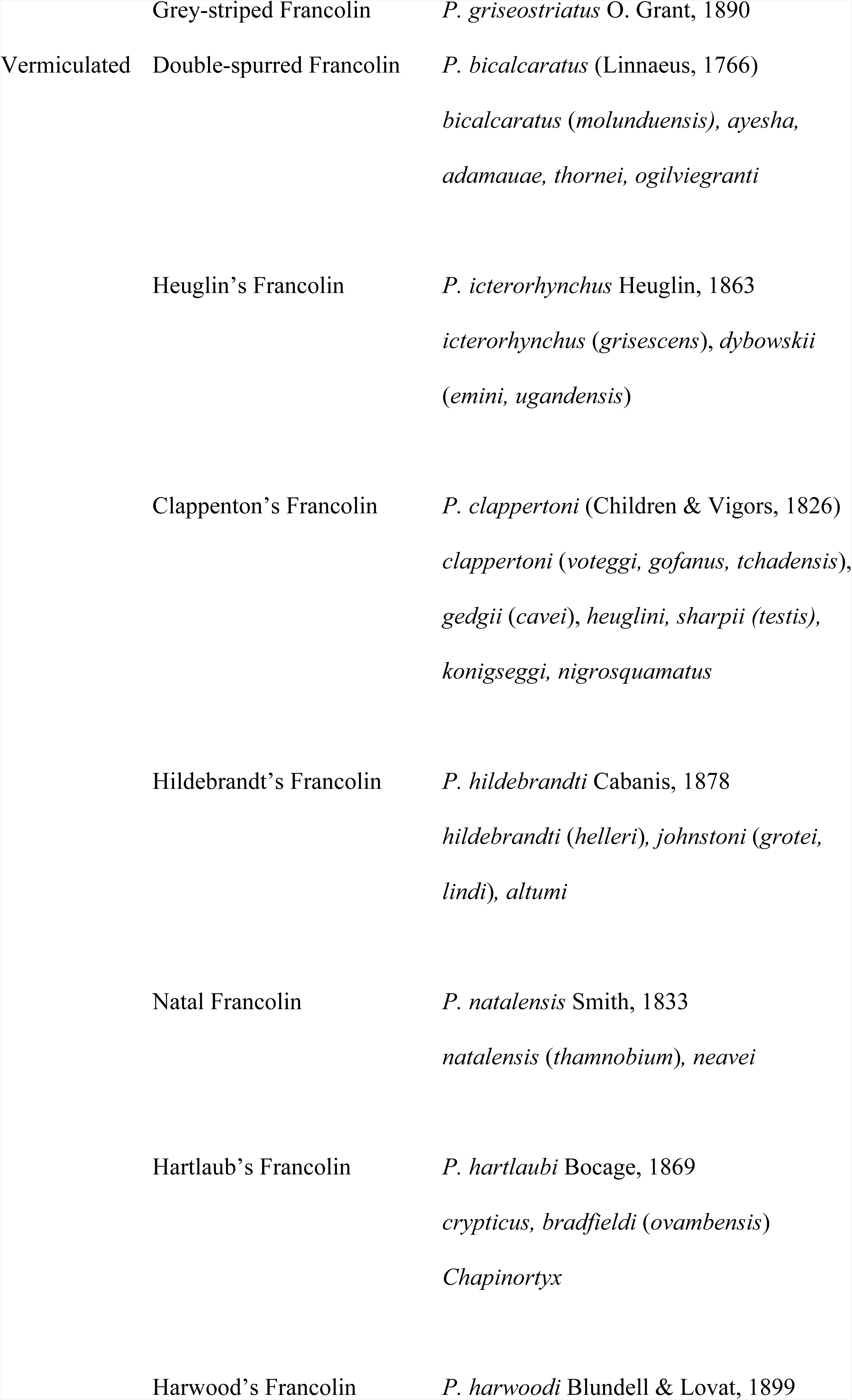
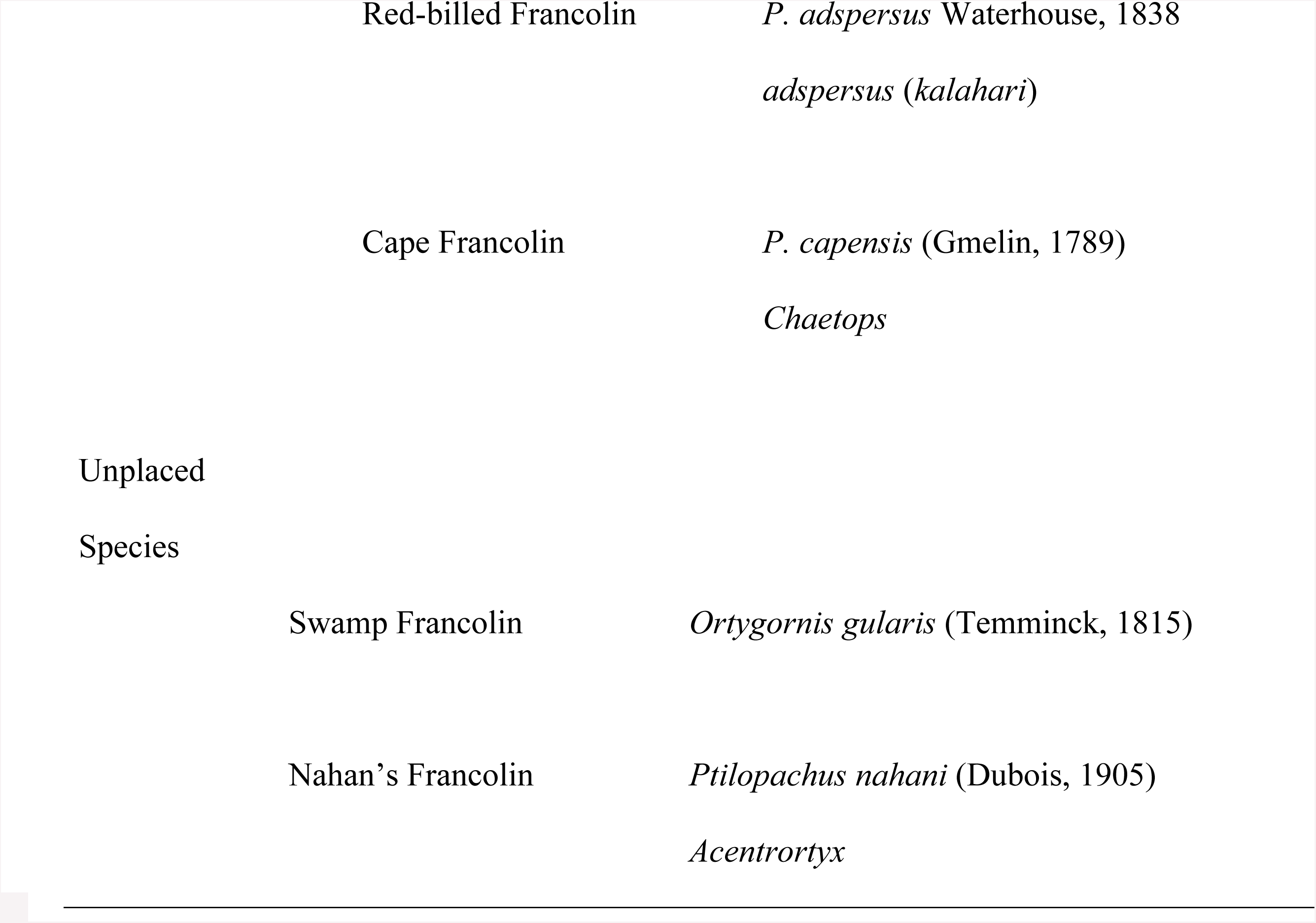
African spurfowl clade, groups, species, subspecies (with synonymized taxa in parentheses) and ‘unplaced’ species recognized by Hall followed by alternative generic epithets [2]. Common and scientific names are according to the IOC list [1].

Hall‘s [2] revision was adopted in many subsequent taxonomic and regional treatments of Galliformes [7, 8, 10, 11]. Nevertheless, other treatments assigned generic status to some of her spurfowl groups and subsets thereof. For example, with regard to African spurfowls, Roberts [12, 13] restricted *Pternistis* to Hall‘s Bare-throated Group, and assigned three of her Vermiculated species (*capensis, natalensis* and *adspersus*) and a fourth (*hartlaubi*) to separate genera, *Chaetops* and *Chapinortyx* respectively. In sharp contrast, Wolters [14] lumped members of her Vermiculated, Montane, Scaly and Bare-throated Groups into a much enlarged *Pternistis*.

Post-Hall’ analyses of francolin/spurfowl syringeal morphology [15], chick plumage [2, 9, 16], vocalizations [17, 18] and DNA [19, 20] decisively reject a sister relationship between Hall‘s two clades of ‘francolins’ [*sensu* 2]. A consensus from the above studies is to recommend phylogenetically placing somewhat modified versions of the Hall francolin and spurfowl taxa within two evolutionarily distantly related phasianine lineages, aligned with the now monophyletic Gallinae and Coturnicinae [20].

For example, the Asiatic Spotted Group (*Francolinus* spp. sensu stricto) and the Swamp Francolin ‘*F.’ gularis* should be removed from the spurfowls and placed within the francolin/*patryse* clade [20], with Spotted/*Francolinus* spp. placed as basal within this clade, and *gularis* with Hall‘s unplaced *’F.’ pondicerianus* with some of Hall‘s Striated taxa (e.g. *sephaena*) [20]. The enlarged ‘true’ francolin assemblage is, in turn, sister to *Gallus* and *Bambusicola* spp. within the Gallinae [20]. Nahan‘s ‘Francolin’ [21] should also be removed from the spurfowl clade and placed outside francolins sensu Hall as sister to another phylogenetically enigmatic African ‘partridge‘, *Ptilopachus petrosus* [22]. These now congeneric *Ptilopachus* spp. (Ptilopachinae) are sister to New World ‘quails’ (Odontophorinae) [23]. African spurfowls (Hall‘s spurfowls minus *Francolinus* spp. sensu stricto, ‘*F*.’ *nahani* and *gularis*) are sister to *Ammoperdix* and *Perdicula* spp., *Coturnix* ‘quails’ and *Alectoris* ‘partridges‘, within the now monophyletic Coturnicinae [20].

In the present study, we deal with the remaining African spurfowls: Hall‘s Montane, Scaly, Vermiculated and Bare-throated Groups [2]. We use more modern species [24] and subspecies [25] concepts and phylogenetic methods to reassess Hall‘s and others’ taxonomic, phylogenetic and biogeographical findings.

## Materials and methods

### Taxon sampling

Taxa and specimens studied herein (Appendix 1) include all putative African spurfowl species, the vast majority of putative subspecies and all specimens examined by Hall [2] at The Natural History Museum (Tring, UK), supplemented by a broader array of material from other major natural history museums mentioned in the Acknowledgements. Where possible, at least 10 specimens for each putative taxon were examined.

### Taxonomy

Taxonomy involves the discovery, description, naming and classification of taxa at all levels of the evolutionary hierarchy. Generally, however, it focuses on the fundamental (terminal) components of biodiversity, which are traditionally species and subspecies [24, 25]. Great emphasis has previously been placed on the relative merits of different ‘species concepts’ with less emphasis on the linked processes through which species are determined [26].

Empirically, we view species as reciprocally monophyletic groups of specimens that are qualitatively similar in terms of suites of diagnostic, consilient characters (from e.g. morphology, vocalizations and DNA markers); and geographically ‘meaningfully’ distributed (e.g. in relation to past/present vegetation types and/or topography, and well-established biogeographical provinces/regions [27]).

Our goal is to identify evolutionarily independent lineages buffered from the homogenizing effects of tokogeny [28]. This is important for spurfowls since hybridization between putative species taxa is thought to be common [29].

Subspecies are groups of populations delineated by geographically steep, congruent clinal variation in multiple characters where their distributional ranges meet. The zones of parapatry (distributional overlap) are characterized by morphologically intermediate individuals or individuals with ‘shuffled‘, undiagnosable sets of characters that appear to reflect hybridization between the largely allopatric populations. Thus, our goal for subspecies is for them to reflect a common phylogeographic genealogy characterized by consilient, potentially adaptive, anatomical, behavioural and ecological differences maintained by constrained interbreeding between taxa. Subspecies in one clade may have geographically similar distributions to those in other clades comprised of much more well-marked evolutionarily significant units [30] and full species. Ultimately, one has to draw the taxonomic line somewhere, with the goal of finding meaningful evolutionary entities. When in doubt, we recognize terminal taxa as subspecies.

A good example of the application of the multifaceted consilient approach is the southern African Black Korhaan (*Eupodotis afra/afraoides*). This taxon was treated as a subspecies pair by avian taxonomists until it was demonstrated that the two taxa were diagnosable as ‘good’ species through a series of consilient evolution of morphological, molecular, vocal, life history and habitat characters; despite evidence of a narrow hybrid zone [31]. Indeed, hybridization, particularly in birds, can still occur even long after speciation [32, 33].

In sharp contrast, 27 putative species/subspecies of Helmeted Guineafowl in the genus *Numida* were combined into a single polytypic species (*meleagris*) with nine subspecies [34]. Subspecies were recognized by high, but imperfect, character consilience, and were connected by narrow to broad zones of ‘hybridization’ between parapatric entities between which there are no discernible differences in courtship behaviour, vocalizations and ecology throughout the entire range of this polytypic species.

Reflecting elements of both the above studies, the sunbird *Cinnyris whytei* of the Malawi highlands was split from the Angolan *C. ludovicensis* [35] due to clear morphological, genetic and distributional differences. But, it was also necessary to describe a newly discovered population of *C. whytei* from Tanzania as a subspecies of *C. whytei* (*C. w. skye*), due to the presence of only minor morphological differences and multi-locus coalescent analyses not being able to exclude the possibility of recurrent gene flow between *whytei* and *skye*. Both examples illustrate the multifaceted consilience approach we adopt, and more generally reflect the view that species are separately evolving metapopulation lineages [26].

In practice within this study, the decision to rank a taxon as a species was made using a consilience framework where entities were:

1. morpho-behaviourally diagnosable (as defined above) and ≥ to 2% difference in unweighted, uncorrected, overall, molecular sequence divergence of mitochondrial DNA [19];
2. reciprocally monophyletic using morpho-behavioural and molecular characters [24]; and
3. were primarily restricted to a commonly accepted biogeographical region or subsection thereof [27].

### Morpho-behavioural characters

The basic body plan of study skins was divided into discrete sections (Fig 1) and scored for variation in colour and patterning: 33 organismal characters reflecting assessment of plumage/integument colour/pattern, measurements of study skins, and vocal characters (Table 2). Measurements (bill length from cere, wing/tail/tarsus/spur length) were taken using a Vernier Calliper or a wing rule.

**Table 2.**
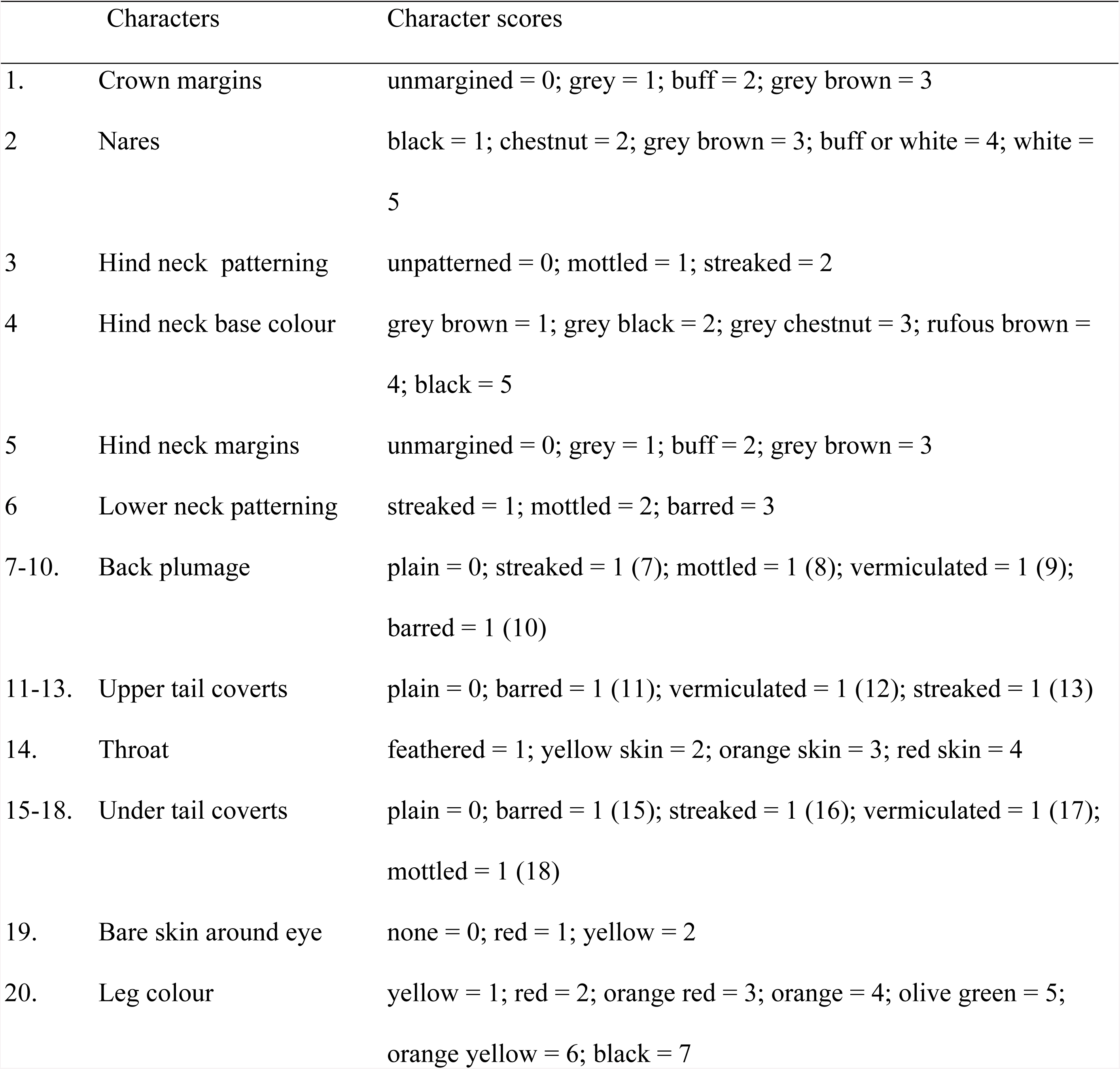
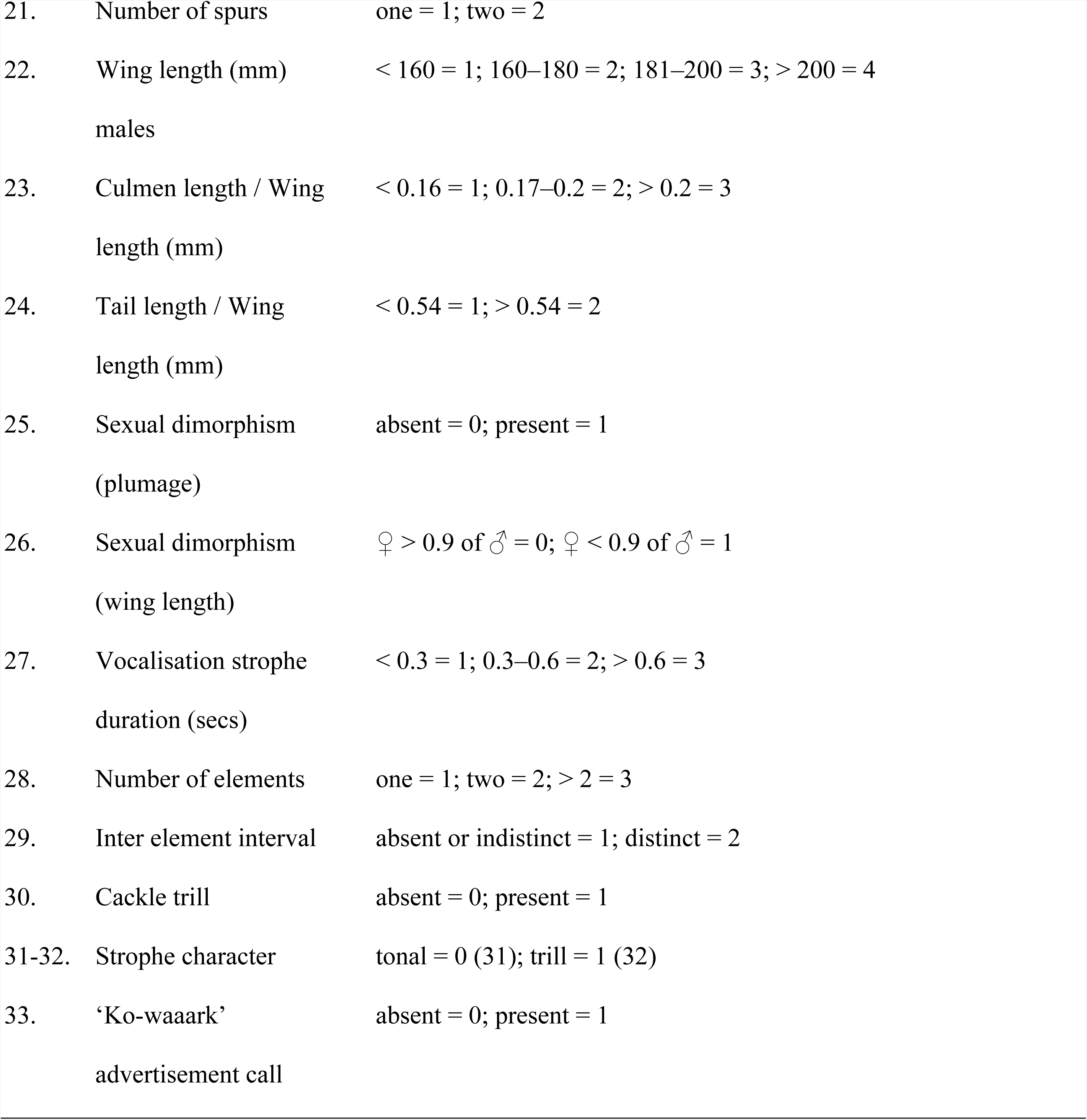
Thirty-three morpho-behavioural characters with scoring criteria used for the phylogenetic analyses of spurfowls.

**Fig 1.**
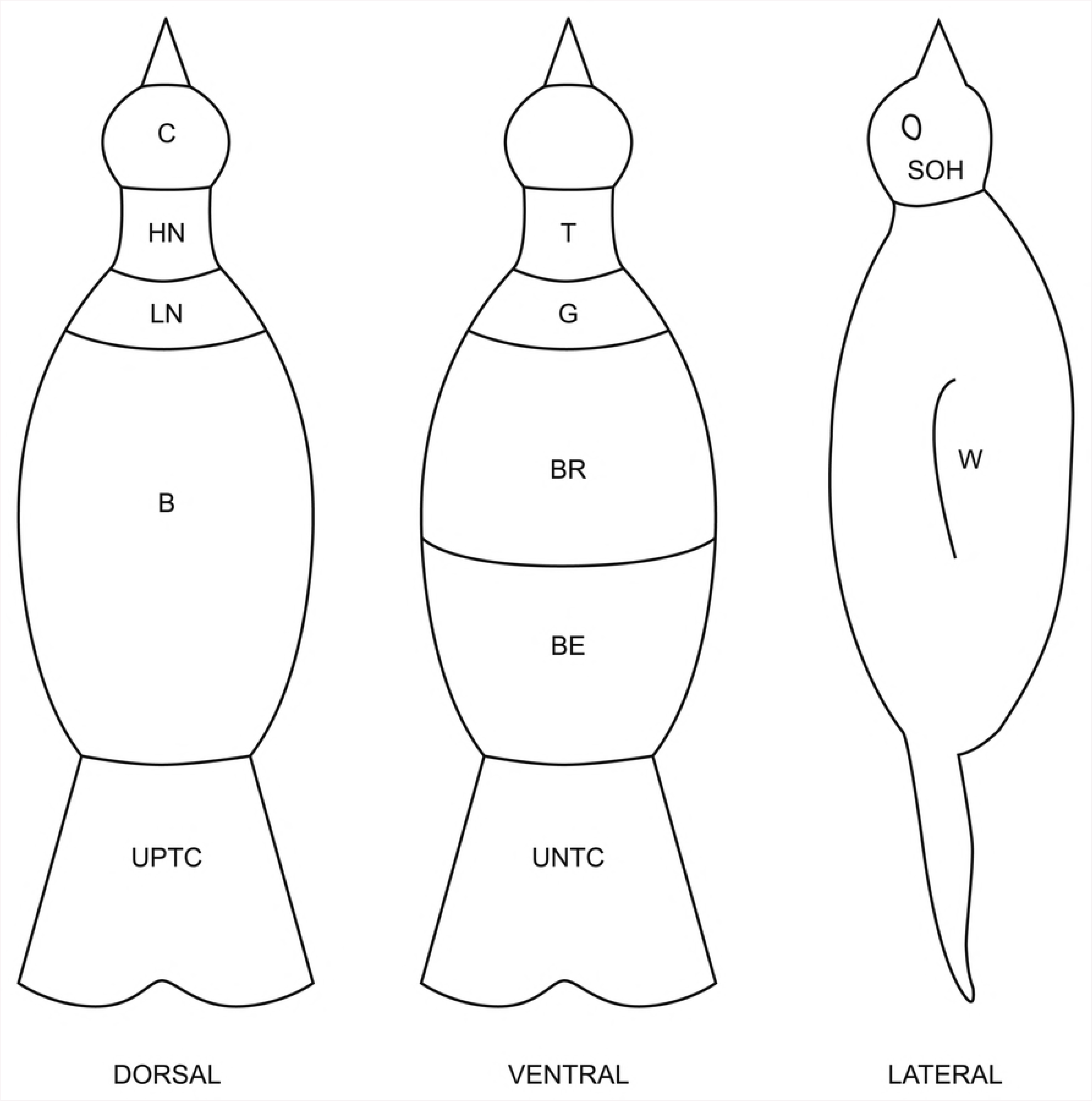
Spurfowl body parts scored when generating plumage characters. C = crown, HN = hind neck, LN = lower neck, B= back, UPTC = upper tail coverts, T = throat, G = throat patch, BR = breast, BE = belly, UNTC = under tail coverts, SOH = side of the head, W = wing.

### Molecular characters and samples

For within-group molecular analyses of spurfowls, 51 putative terminal taxa were studied (Table 3) with respect to four mitochondrial markers: Cytochrome *b* (CYTB - 1143 base pairs), Control region (CR - 820 bp), NADH dehydrogenase subunit 2 (ND2 - 1041 bp) and 12S rRNA (12S - 706 bp); three nuclear DNA markers: Ovomucoid G (OVO-G - 449 bp), Glyceraldehyde-3-phosphodehydrogenase (GAPDH – 361 bp) and Trans Globulin Growth Factor Beta2 intron-5 (TGFB - 596 bp) (Appendix 1).

**Table 3.**
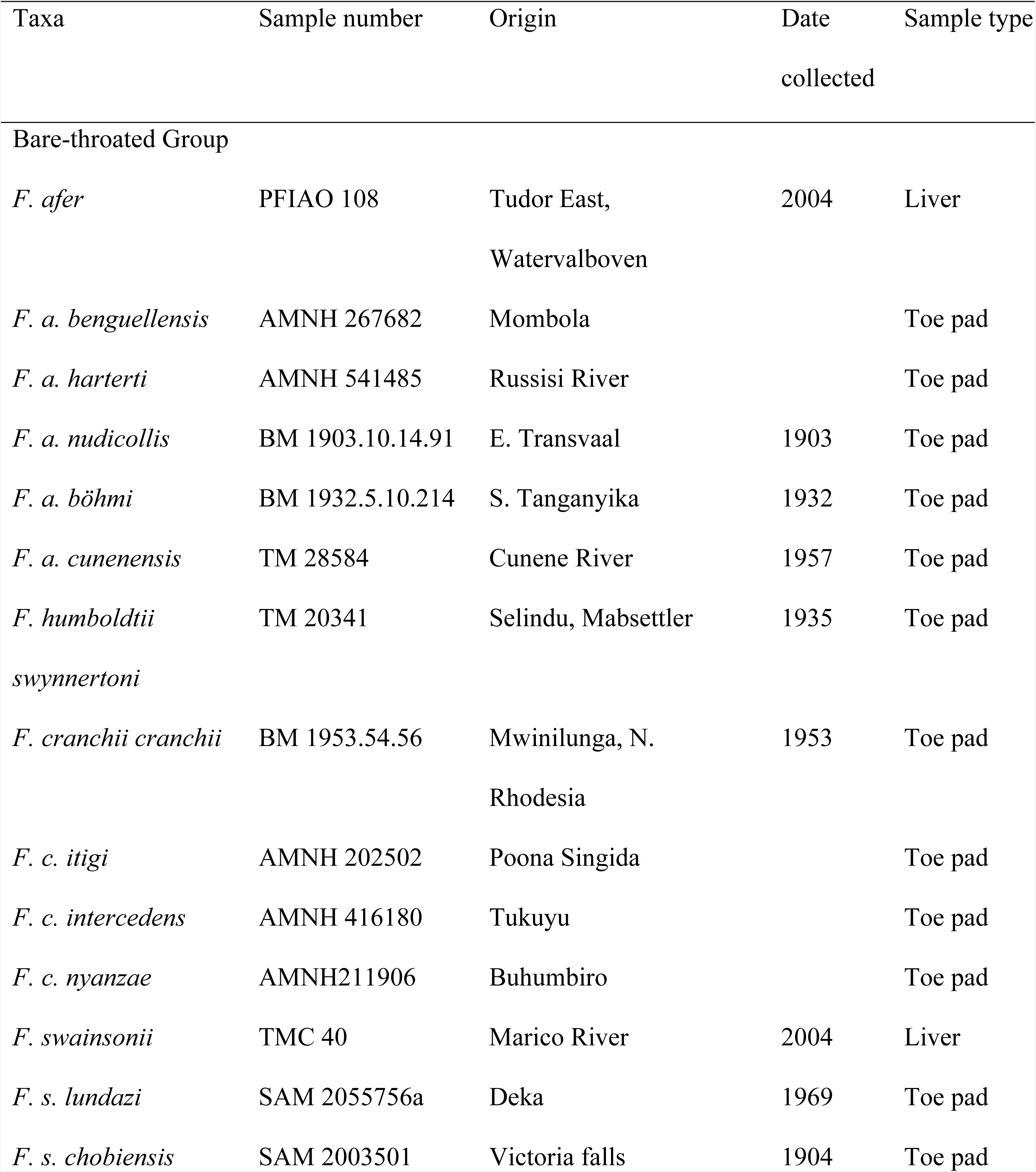
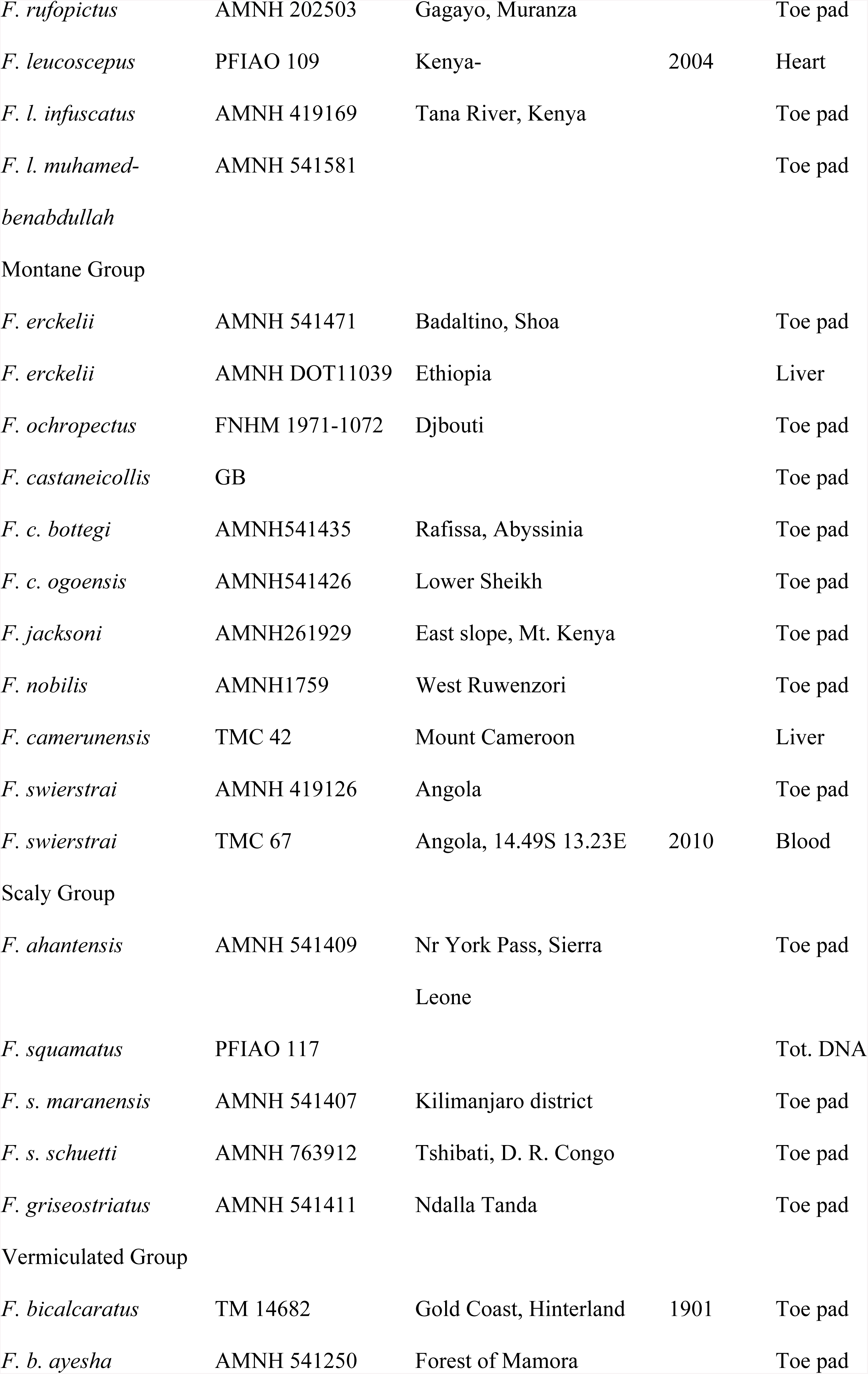
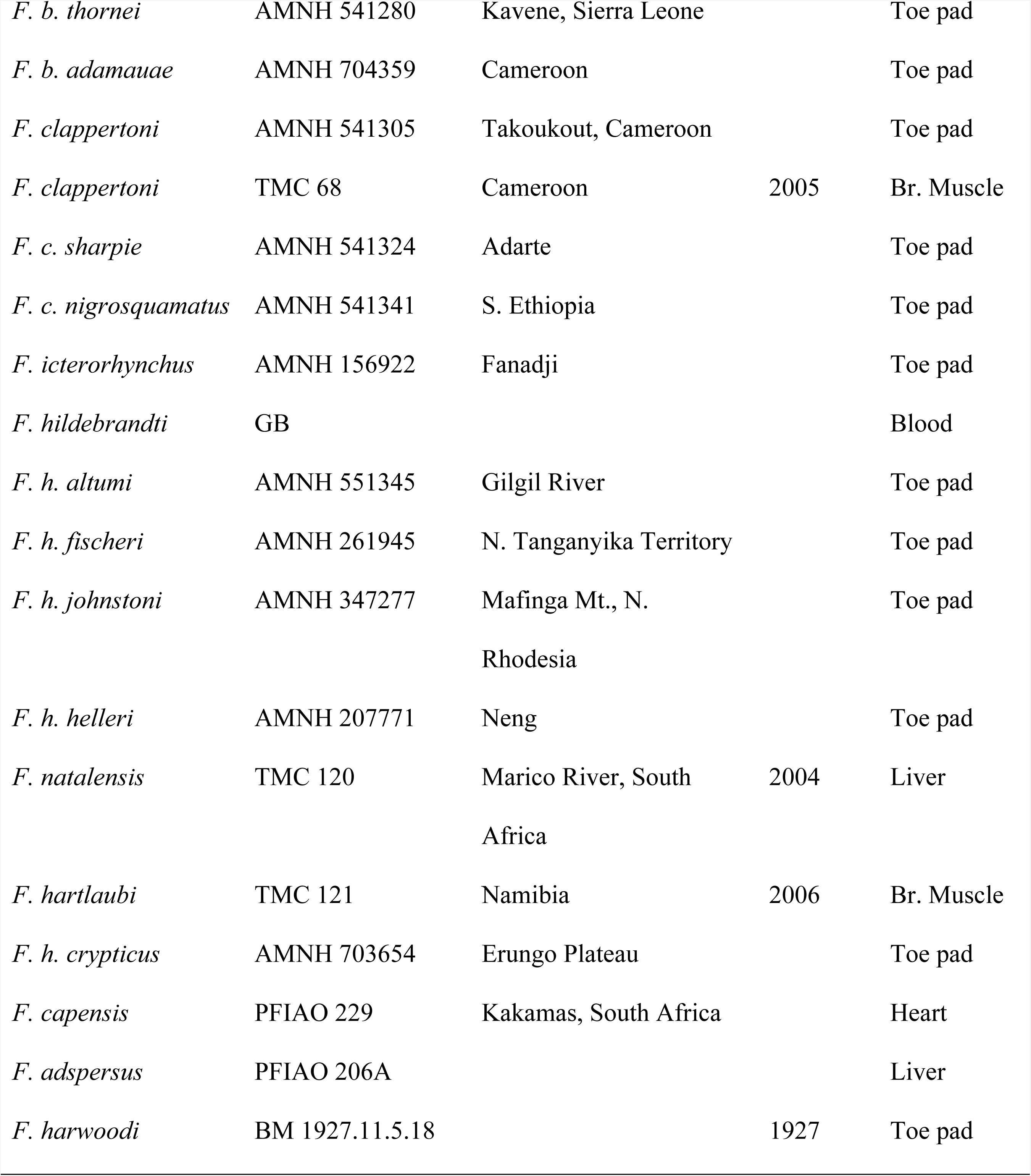
Sample information for spurfowl taxa recognized by Hall [2] for which DNA sequences were generated. Acronyms; AMNH = American Museum of Natural History, FHHM = French Natural History Museum, TM = Transvaal Museum - Ditsong National Museum of Natural History, BM = British Museum - Natural History Museum at Tring, SAM = Iziko Museums of Cape Town (Natural History), FIAO = FitzPatrick Institute of African Ornithology, TMC = Timothy M. Crowe, University of Cape Town, South Africa, GB = GenBank, Br. muscle = Breast muscle. Generic terminology follows that generated in this study.

Primers used in sequencing are listed in Tables 4 and 5. The 1143 bp long CYTB was sequenced for all taxa included in this study while data for the other markers may be missing for some taxa. Contrary to earlier work [19, 36] which focused on few species, all putative species and most subspecies attributed to African spurfowls were included (Appendix 1). Some 72% of specimens sequenced in this study derived from DNA extractions of toe-pad scrapes off museum skins. As a result, only CYTB was sequenced for both fresh and historical tissues and the other six markers were sequenced for species for which there were fresh tissues. Due to the fragmented nature of the historical sourced DNA, the CYTB gene for the toe-pads was sequenced in multiple fragments (six for each sample) using spurfowl-specific primers (Table 5).

**Table 4.**
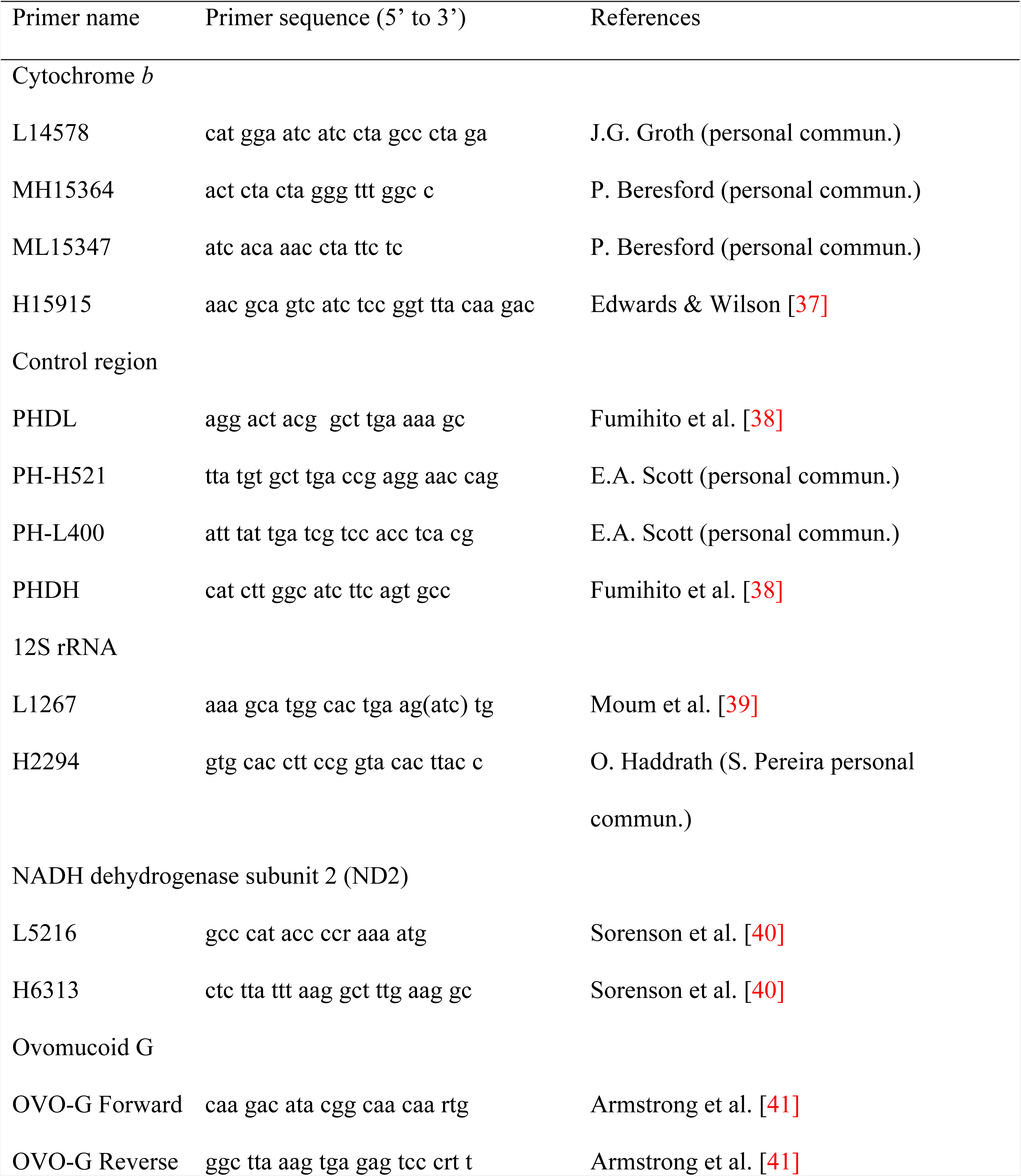
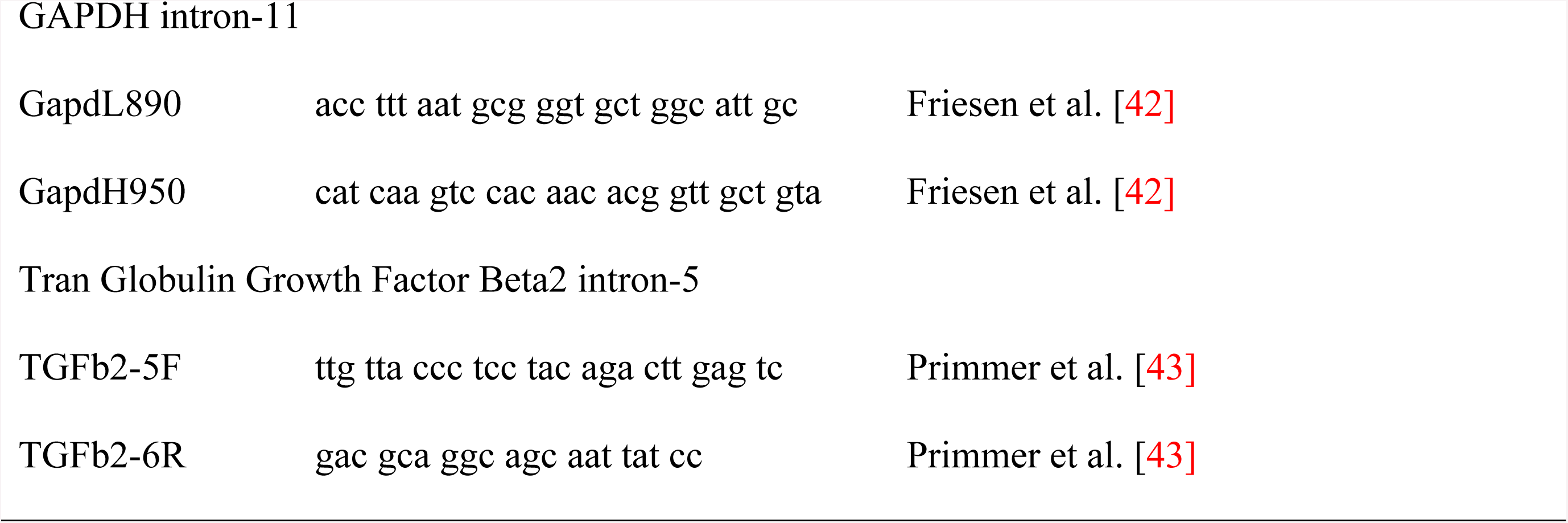
DNA markers sequenced and primers used for PCR-amplification and sequencing of preserved tissues.

**Table 5.**
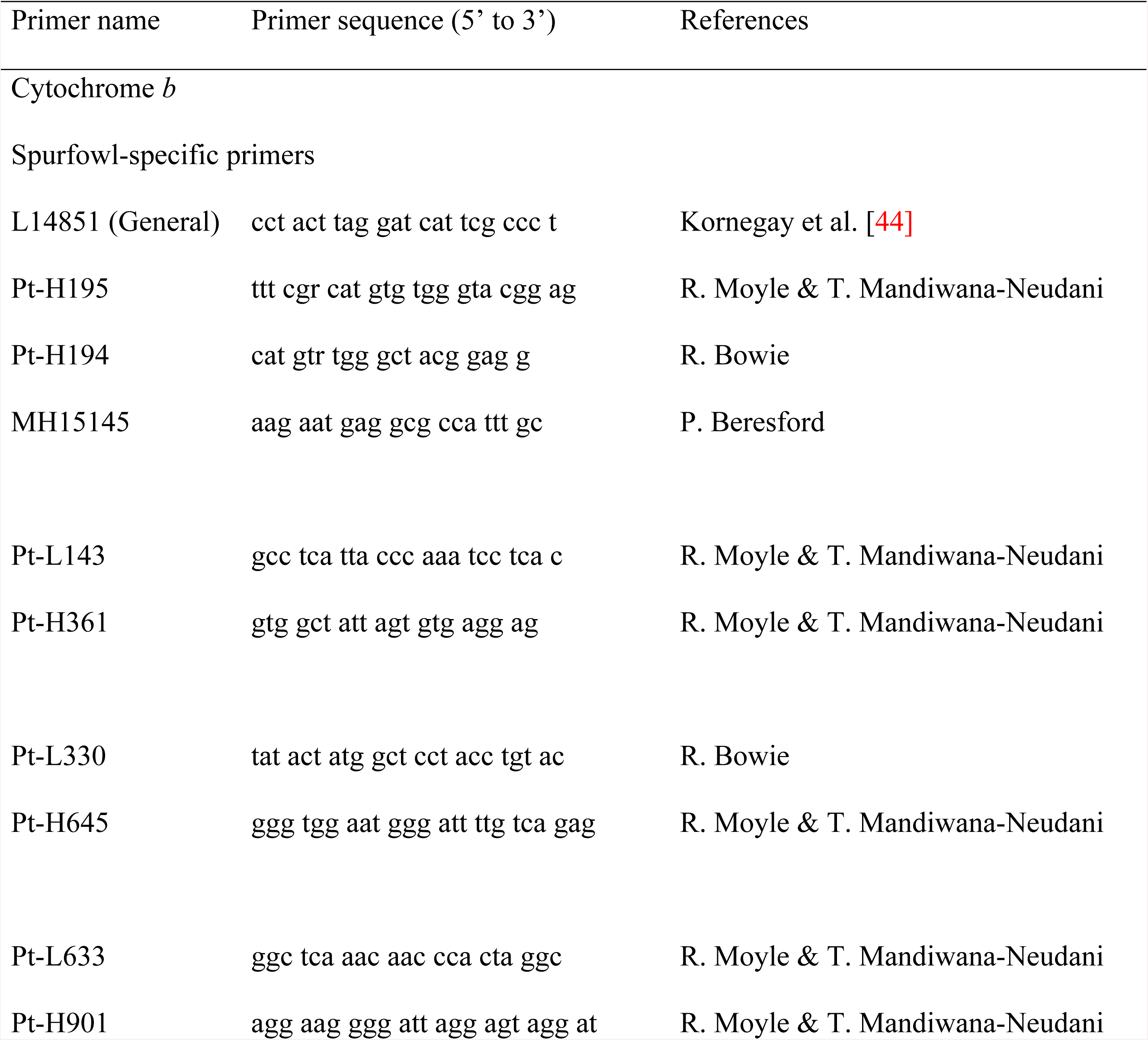
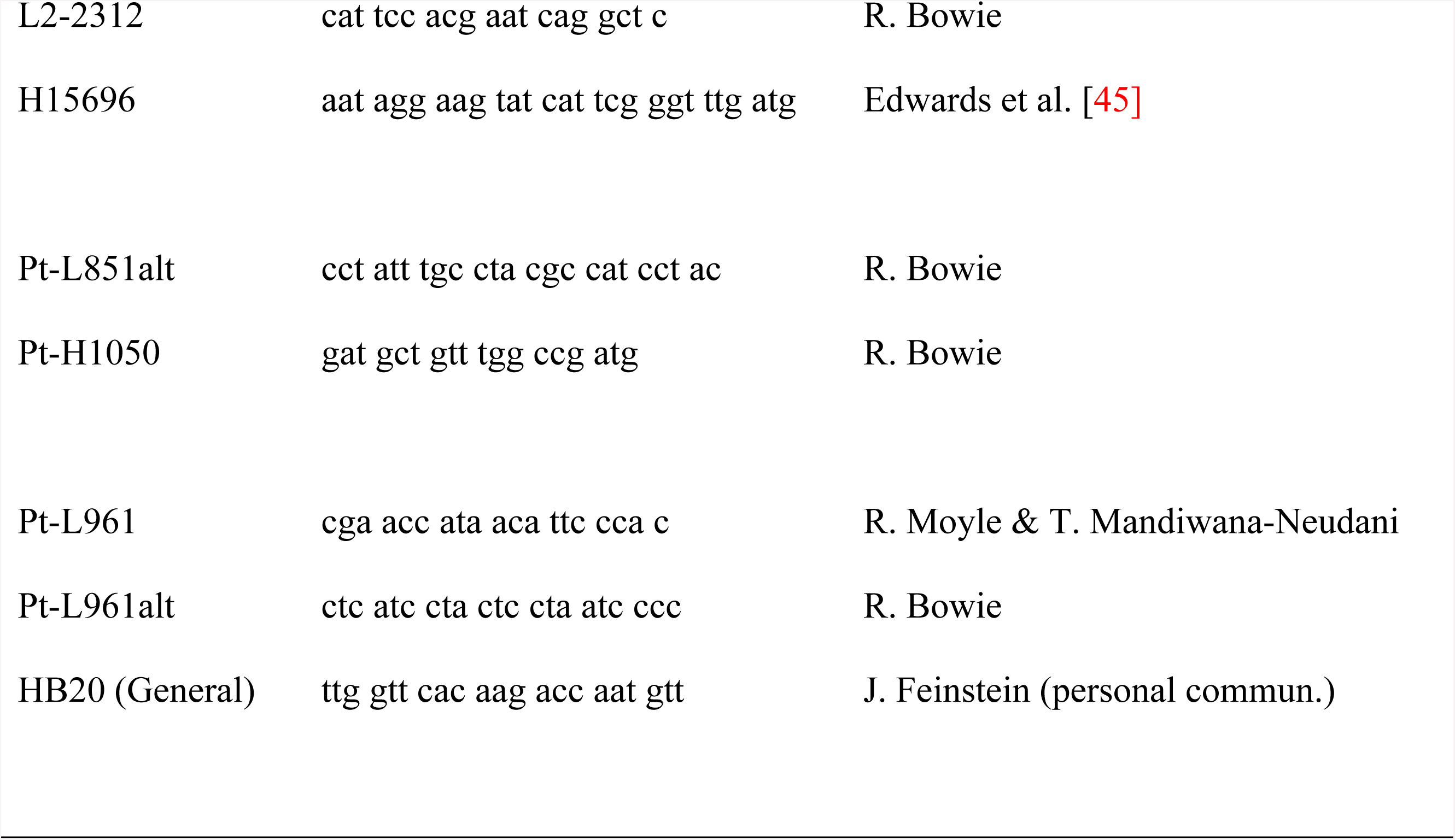
DNA markers sequenced and primers used for PCR-amplification and sequencing of museum toe pads.

### Phylogenetic methods

Taxa were placed phylogenetically, following the principle of character consilience to reflect progressively more inclusive reciprocally monophyletic groupings [46]. Qualitative morpho-behavioural characters (morphology, behaviour, life history) were analyzed in combination with DNA sequence characters, in a ‘total evidence’ phylogenetic analysis. This approach was chosen because combined data sets may show clade support and resolution that is ‘hidden’ by separate analysis of character partitions. For instance, when data are concatenated, different types of characters that evolve at somewhat different rates may ‘click in’ at different levels of phylogeny [i.e. deep, shallow and intermediate nodes; 47, 48].

Parsimony was employed as the optimality criterion for the combined DNA and morpho-behavioural character analyses [49]. Indeed, the meta-analysis of more than 500 articles using model- and parsimony-based methods found strongly supported topological incongruence in only two of the studies examined [50].

All the data matrices were rooted on *Perdicula asiatica* and *Ammoperdix heyi* following [4]. For inter-taxon genetic distances, uncorrected pairwise distances were calculated in PAUP ver. 4.0b10 and were transformed into percentages.

### Distributional range maps

Another challenging and indispensable aspect in the analyses outlined below was to produce maps showing the distributional ranges of the various spurfowl taxa ultimately recognized. In Step 1 in developing the range (as opposed to point locality) maps for each taxon that emerged, the ‘Atlas of Speciation in African Non-passerine Birds’ [51] was used since it still presents the best distribution ranges of species produced from the point localities of the specimens collected. This was supplemented in Step 2 - consulting the ‘Atlas of Southern African Birds’ [52] which was helpful in filling distribution gaps for southern African species. Step 3 involved using Hall‘s inferred distributions [2] to complete the ranges of species and subspecies recognized.

## Results

### Morpho-behavioural characters

Character information for morpho-behavioural characters are presented in Table 6.

**Table 6.**
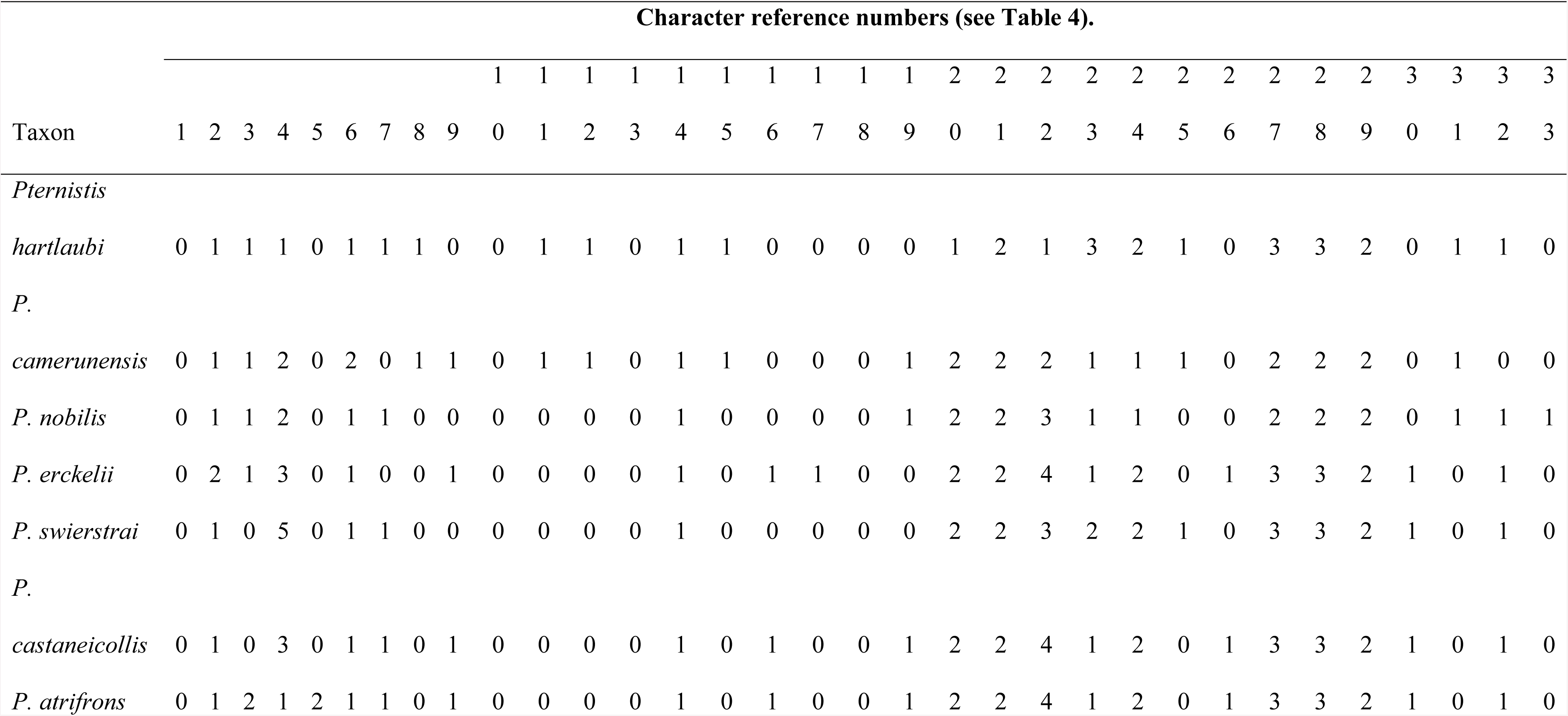
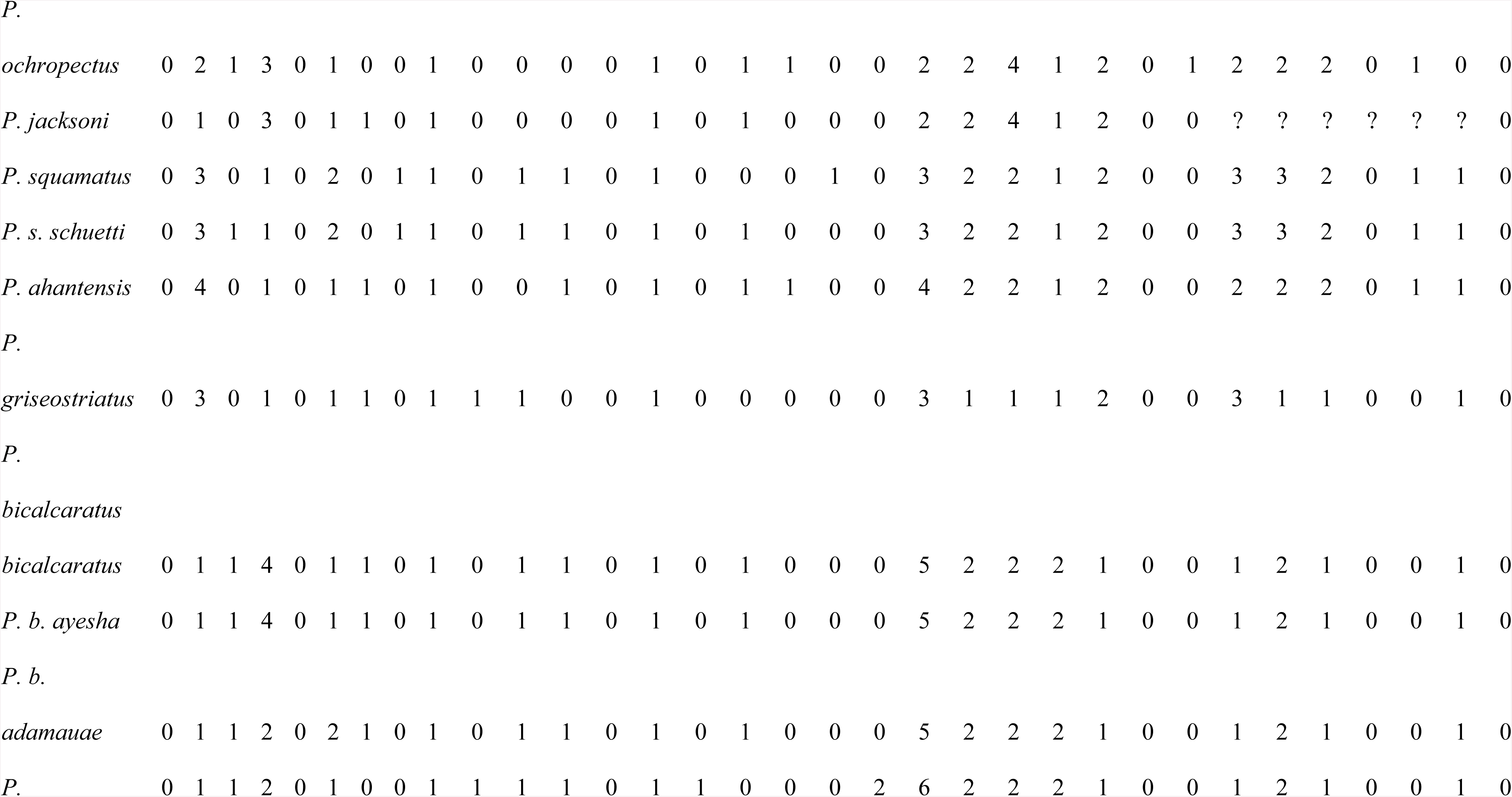
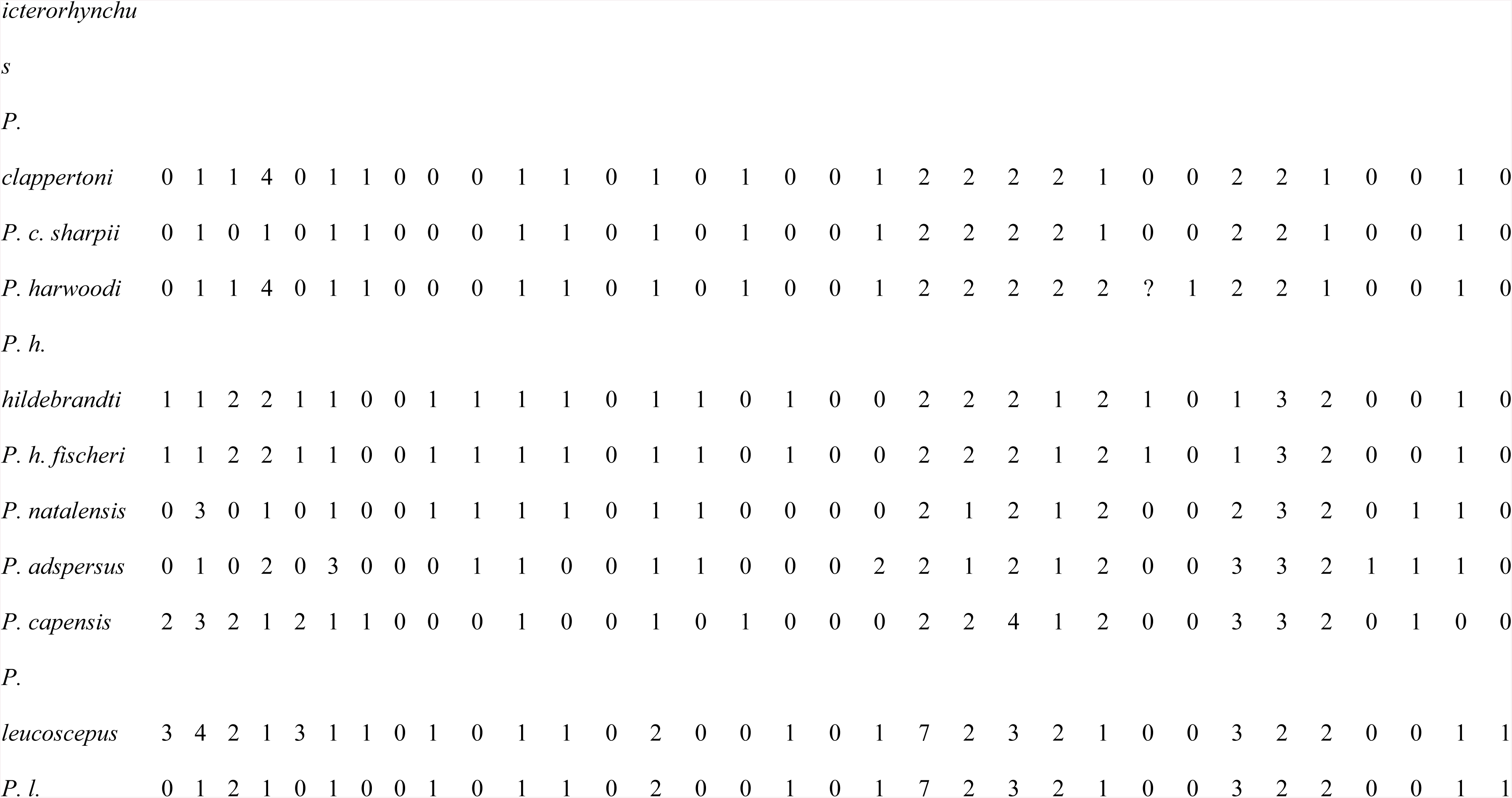
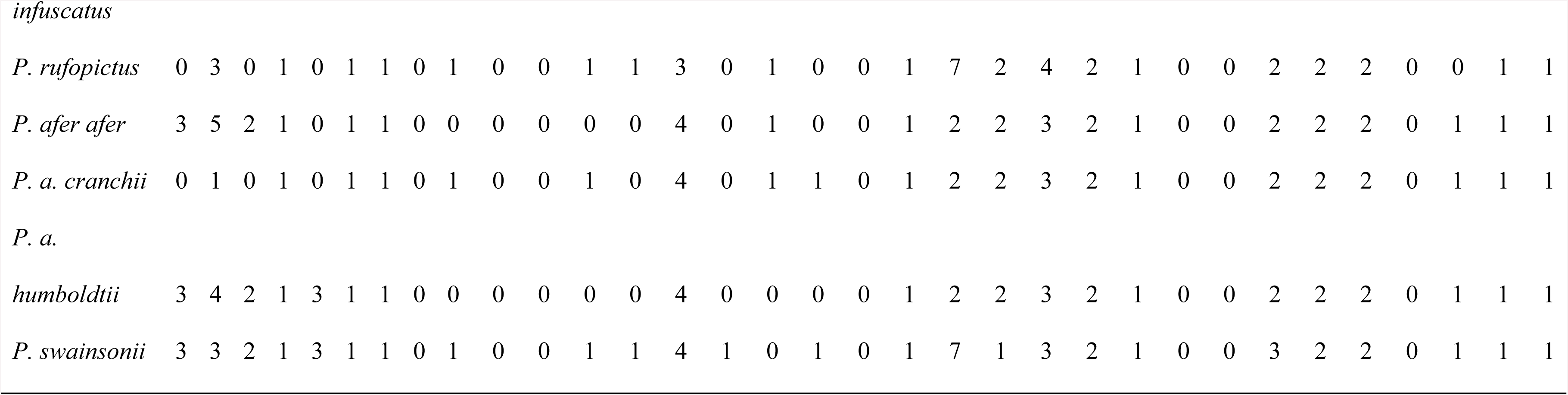
Morpho-behavioural character scores matrix used for the phylogenetic analysis of spurfowls.

### Phylogenetics

The ‘total evidence’ parsimony analysis based on 5149 characters (33 organismal and 5116 DNA bases) and 33 terminal taxa produced two equally parsimonious trees of length 2124, the strict consensus of which is presented as (Fig 2). Since only one of Hall‘s spurfowl species groups [2], the phylogenetically terminal Bare-throated Group, emerged as monophyletic and the others are para- or polyphyletic, we recognize only one monophyletic genus for the African spurfowls: *Pternistis.*

**Fig 2.**
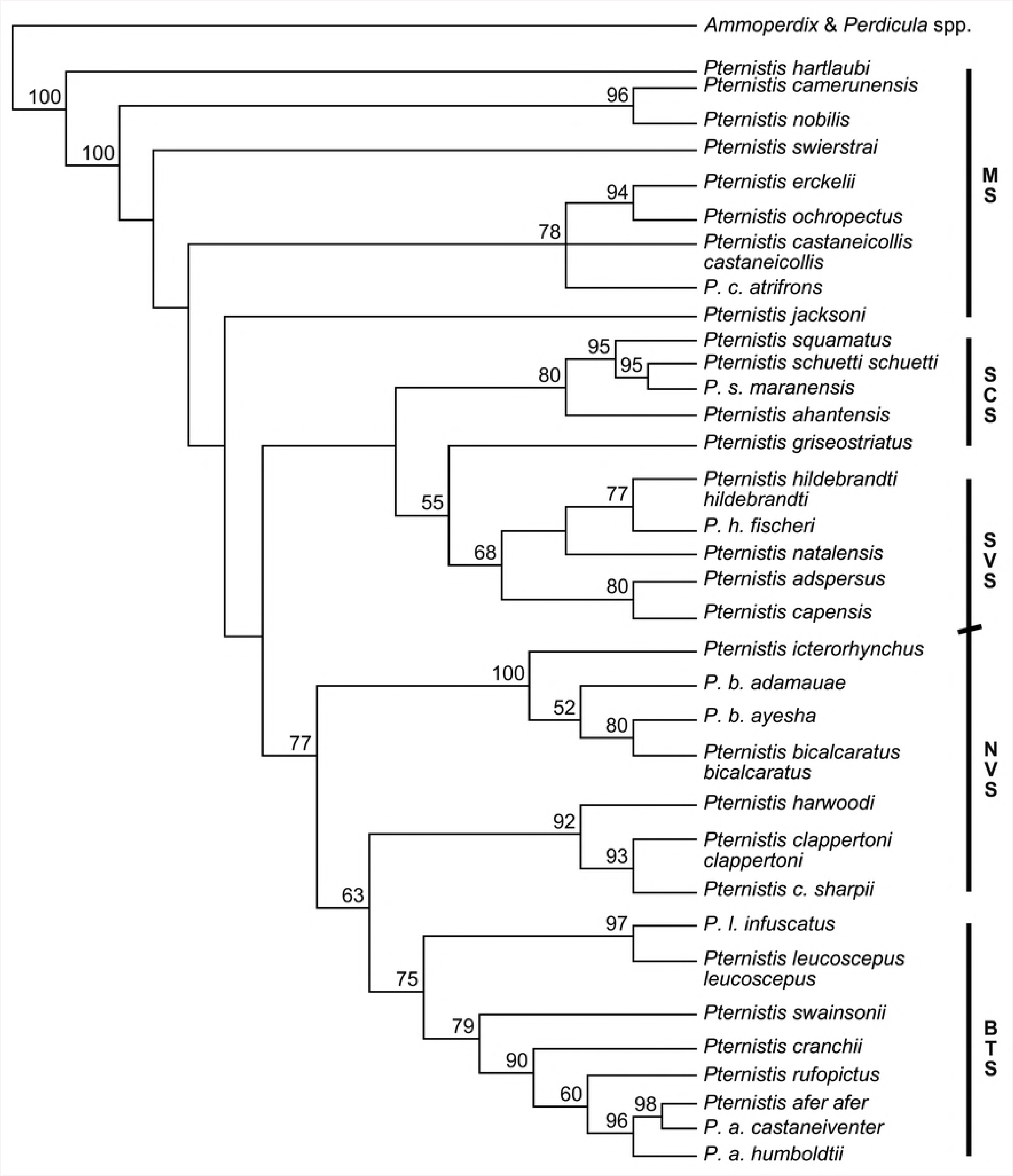
Strict consensus parsimony tree for spurfowls constructed from two most parsimonious trees. Numbers mapped above nodes are jackknife support values. MS = Montane spurfowls, SCS = Scaly spurfowls, SVS = Southern Vermiculated spurfowls, NVS = Northern Vermiculated spurfowls and BTS = Bare-throated spurfowls.

### Cladogenesis

*Pternistis hartlaubi*, one of Hall‘s Vermiculated taxa [2], is the basal African spurfowl. Hartlaub‘s Spurfowl occupies dense, mixed grass-shrub cover on boulder-strewn slopes and rocky outcrops in hilly and mountainous regions within a granite and sandstone substrate surrounded by semi-desert open savanna [53]. It is confined to central and northern Namibia, particularly on the Namibian escarpment and extreme southwestern Angola [54].

The upper mandible of *P. hartlaubi* is horn coloured and the lower yellowish. The male has a dark grey-brownish crown, a pronounced white eyestripe, offset by a black line below and chestnut ear coverts. The back is grey, faintly streaked and barred with brown. The belly is pale grey, heavily streaked with brown. The black and white under-tail coverts are conspicuous in flight and in courtship display. The adult female has an orange-brown eyestripe, and a grey-brown head, cheeks, chin and belly. The back is grey-brown with strong vermiculations [53].

Hartlaub‘s Spurfowl is markedly distinct from other African spurfowls [2, 53]. Indeed, it differs from ‘francolins’ *sensu lato* in general, in that it: (1) has markedly sexually dimorphic plumage [2]; (2) has a disproportionately long bill used for digging underground corms and tubers [53]; (3) is the smallest spurfowl and is markedly sexually size-dimorphic (males 245-290 g., females 210-240 g. – 54]; (4) has yellow (normally black or red/orange-red in spurfowl) tarsi with virtually no tarsal spurs – actually only tiny bumps [2]; (5) is socially monogamous throughout the year [53]; (6) has vocalizations markedly different from (but still link with) the rest of the spurfowls [17, 18]; (7) demarcates and defends its territory year-round, using a combination of uniquely antiphonal duet calling (initiated by the hen) and displays, rather than overt aggression [53]; and (8) seems not to require standing/flowing water for drinking [53].

With regard to putative subspecies, populations from southern Angola (nominate ‘*hartlaubi*‘) are somewhat smaller than those from Namibia. Those from the Kaokoveld and Erongo (’*crypticus*‘) are paler than those from the Waterburg and Otavi (’*bradfieldi*‘) in the east. We regard these differences as clinal variation. The two specimens (from Erongo and the Waterburg) were 0.4% CYTB divergent. We recognize no subspecies for this taxon.

Hartlaub‘s Spurfowl‘s closest CYTB taxon is *P. squamatus* at 7.8% sequence divergence.

Hall‘s [2] Montane spurfowls follow on phylogenetically from *hartlaubi*, but are paraphyletic (Fig.2). They are forest-dwelling taxa, forming two, monophyletic clades: *camerunensis + nobilis* and *erckelii + ochropectus + castaneicollis*, linked by *swierstrai*. Then comes *jacksoni* as a link to the also paraphyletic, lowland/secondary forest-dwelling Scaly spurfowls.

Thereafter come the also paraphyletic woodland, savanna, scrub and bush dwelling Vermiculated (divided into northern and southern assemblages) and the monophyletic Bare-throated taxa.

### The Montane spurfowls

There are seven Montane spurfowl species, one with two subspecies: *swierstrai, camerunensis, nobilis, erckelii, ochropectus, castaneicollis* (*castaneicollis, atrifrons*) and *jacksoni.*

They are distributed across the mountains of north-eastern Africa from Eritrea to Mt. Kenya, extending west through the Albertine Rift, to Mt. Cameroon and south to the highlands of Angola [2] (Fig 3). Montane spurfowls are confined to forested habitat, which provides roosts and cover, although some taxa (e.g. *P. erckelii*) will venture out into wooded scrub, heath and grassland with shrubs [54].

**Fig 3.**
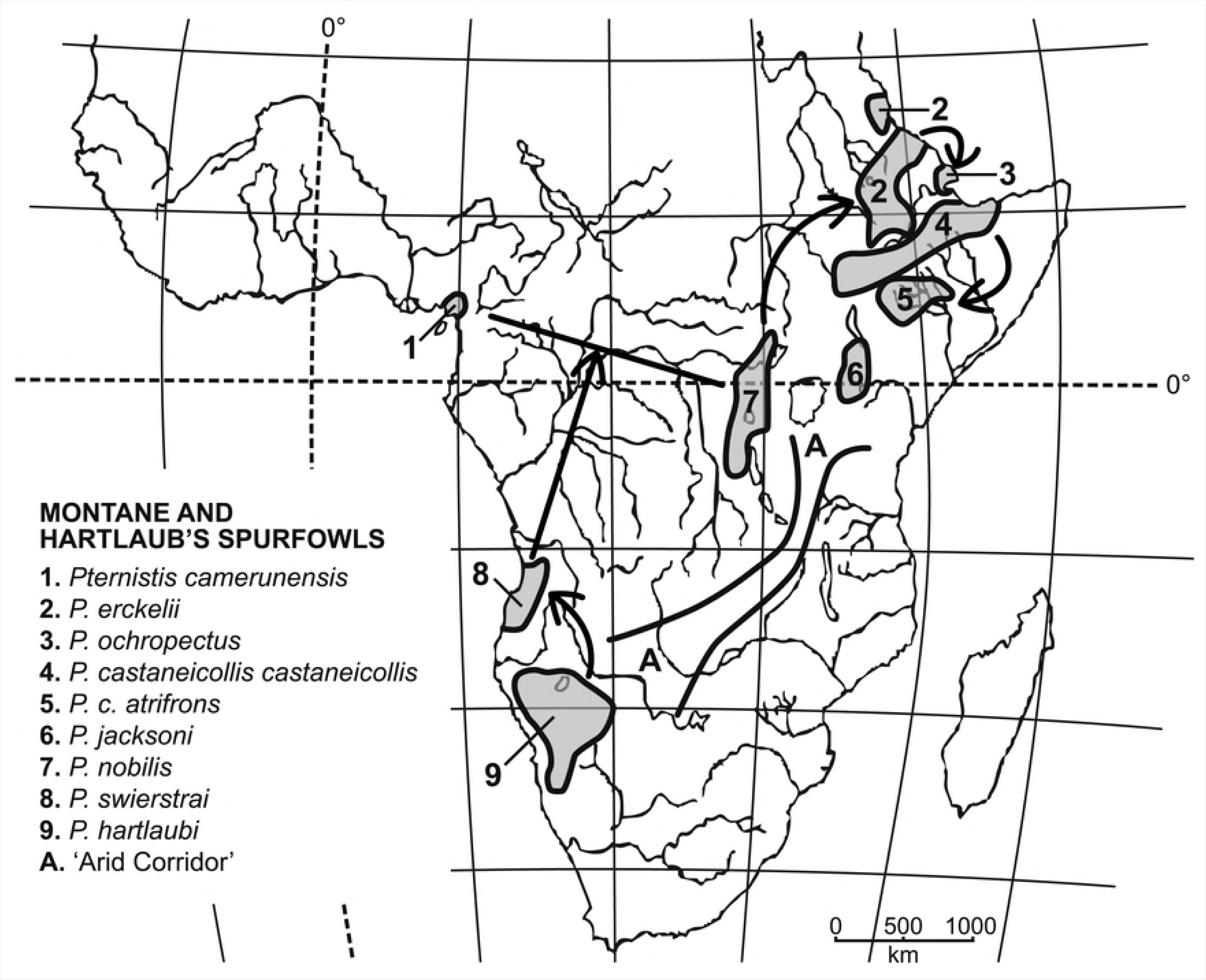
Geographical distributions of Hartlaub‘s Spurfowl, Montane spurfowls and the ‘Arid Corridor‘. Arrows draw attention to phylogenetically sequential cladogenesis.

The Montane spurfowls are the morphologically least homogeneous of Hall‘s spurfowls [2]. There is no diagnostic ‘Group’ morphological character other than that the males have the crown, lower back, primaries and tail plain brown or red-brown. Females of the relatively small, moderately sexually dimorphic species (*P. camerunensis, P. swierstrai)* have vermiculated primaries, lower back and tail.

Variation in some characters follows geographically clinal trends, with the birds of the extreme northeast being: the largest and most heavily spurred with dark bills, yellowish tarsi, no bare skin round the eyes, with the sexes alike [2]. The two isolated, sexually dimorphic western species (*camerunensis* and *swierstrai*) are the least heavily spurred and the smallest species. Thus, they most closely resemble the basal, and also sexually dimorphic, *P. hartlaubi*. The central African species, *nobilis*, ‘connecting’ these three species to those in the northeast is of intermediate body mass [54]. Generally, Montane spurfowls differ from one another primarily in their belly plumage, particularly on mid- and lower belly.

The closest non-montane CYTB taxon to them is *P. squamatus* at 5.3% sequence divergence.

*Pternistis swierstrai* is an uncommon, endemic of Angola, confined to undergrowth within patches and edges of relict evergreen forest in the highlands of western Angola, Mountains Moco and Soque, the Bailundu highlands and Mombolo Plateau along the escarpment, with isolates on the Chela escarpment, Tundavala (Huila District) and Cariango (Cuanza Sul District) (Fig 3) [54]. It ventures into grass- and bracken-covered slopes and gullies [54].

Swierstra‘s Spurfowl is a small spurfowl (both sexes 375–565 g [54]), and has an orange-red bill, a yellow ear-patch, yellow eye-ring on males (blue in females), red tarsi with one spur, only in males. It is weakly sexually dimorphic in plumage. Both sexes have a conspicuous white eyestripe and throat, brown back plumage (irregularly blotched in the female). The male‘s black breast contrasts with the white throat, whereas the lower belly feathers have broad buff central streaks with blackish margins. The belly plumage of the female is white, barred/blotched with dark brown [2].

Its closest CYTB taxon is *P. squamatus* at 5.3% sequence divergence.

*Pternistis camerunensis* is sister to *nobilis.* It is endemic to, and locally distributed within dense undergrowth and edges of forests on the south-eastern slopes of Mt. Cameroon, between 850 and 2100 m above sea level (Fig 3) [54]. The Mt Cameroon Spurfowl is a small (male ∼593 g., female ∼509 g.), sexually dimorphic spurfowl, and has an orange-red bill, red eye-ring, and orange-red tarsi with 1-2 spurs, only in males. The male has a dark brown crown and nape. Its throat is grey-buff with the belly feathers chestnut with grey edges. Its upper tail coverts and primaries are grey brown, and wing coverts and the lower neck are deep maroon, with light grey scalloping on the lower neck. Its back is rich dark brown (excluding the lower neck). The belly and lower neck are plain grey with some black feather centres and shaft streaks. The chest and belly plumage of the female is mottled and vermiculated with black, dark brown and buff with some off-white U-to V-patterning on the belly and lower neck.

The closest CYTB taxon to *P. camerunensis* is its sister-taxon, *P. nobilis,* at 7.4% sequence divergence.

*Pternistis nobilis* [54] is endemic to the highland Ruwenzori and Kivu forests in the Albertine Rift and mountains in far western DR Congo, south-western Uganda and borders of Rwanda and Burundi, and is locally common in dense undergrowth, forest edge and moist bamboo thickets (Fig 3). The Noble Spurfowl [54] is medium-sized and sexually monomorphic (males averaging 877 g., females 635 g.]. It has a red bill, eye-ring and tarsi with 1-2 spurs (upper shorter), only in the male. It has a grey-brown head, primaries and rump, and a buff throat. It is dark maroon overall, particularly on the wings and back, with light grey scalloping on the lower neck. The rest of the belly feathers are chestnut with narrow grey or whitish edges or scallops [2].

With regard to subspecies of *P. nobilis*, we regard ‘*chapini*‘, from the Ruwenzori Mountains as an idiosyncratic variant since it differs only by having somewhat narrower greyish edges to the belly feathers [2].

The closest CYTB taxon to *P. nobilis* is its sister-taxon, *P. camerunensis,* at 7.4% sequence divergence.

*Pternistis erckelii*, the most northerly distributed Montane spurfowl, is sister to *P. ochropectus*. It is distributed in giant heath, forest scrub remnants and edges above 2000 m, extending, relatively continuously up to 3000 m, from the vicinity of Addas Ababa in the massif of central and northern Ethiopia southwards to southern Eritrea (Fig 3). Unlike other Montane spurfowls, it will venture out of forest into adjacent heath and grassland. Erckel‘s Spurfowl is the largest African spurfowl (males 1050-1590 g., one female 1136 g. [54]). It has a black bill and yellowish tarsi with two spurs (upper longer), only in the male. It is sexually monomorphic, and has a black forehead and eyestripe, chestnut crown, grey ear coverts and white throat. Its lower neck is grey like the upper belly, but with greyish brown margins and a thin central buff streak, whereas the upper belly feathers have central greyish black streaks. Lower belly feathers have a broad buff central streak constricted in the middle and expanded distally into a tear-drop, margined with rufous [2]. The “somewhat greyer” [2] putative subspecies, ‘pentoni‘, an isolated population from the Red Sea Hills at Erkowit, is not recognized.

The sister and CYTB closest species to *P. erckelii* is *P. ochropectus* at 2.6% sequence divergence.

*Pternistis ochropectus* is a large spurfowl (one male 809 g., one female 605 g. [54]) endemic to the evergreen juniper forest mostly above 1200 m. on the Plateau du Day of Djibouti (Fig 3). The Djibouti Spurfowl [54] has a black bill with the lower mandible yellowish and yellow tarsi with two spurs (upper longer), only in the male. The lower belly feathers of *P. ochropectus, P. erckelii* and *P. castaneicollis* are similar, but *P. erckelii* and *P. castaneicollis* are more heavily marked with brown on the back and breast. The belly feathers of *ochropectus* have a broad buff central streak constricted in the middle and expanded distally into a tear-drop, margined by a greyish black U-shaped streak [2].

The CYTB closest and sister-species to *P. ochropectus* is *P. erckelii* at 2.6% sequence divergence.

*Pternistis castaneicollis* is a large spurfowl (males 915-1200 g, females 550-650 g. [54]), and is restricted to montane heath moorlands, juniper forests and forest edge/scrub above 2800 m. It extends broadly in montane ‘islands’ along the mountain ranges of central and south Ethiopia on both sides of the Rift Valley to Somalia in the extreme northwest, and to the Kenyan border in the extreme south (Fig 3) [54]. The Chestnut-naped Spurfowl is morphologically geographically variable, but most similar to *P. erckelii* [54]. It has a red bill, yellow ear-patch, yellowish eye-ring in males (blue in the female) and orange-red legs with two equally long spurs, only in the male. It is sexually monomorphic in plumage, but females are smaller. It has less black on the face than *erckelii* and *ochropectus*. Its belly feathers having a broad buff central streak, constricted in the middle and expanded distally into a tear-drop, margined with rufous. Its eastern Ethiopian populations have an extensive double-U-patterning on the back with wing coverts and breast clearly defined in black and white, with some ochre and chestnut, grading to mainly white on the belly [2].

The closest CYTB taxon is *P. erckelii* at 4.2% sequence divergence.

The subspecies *atrifrons* (for which we had no DNA sequence data) is confined to the Mega Mountains of southern Ethiopia (Fig. 3). It was recently elevated to full species [55] and is 1.2-1.3% CYTB divergent from *P. c. castaneicollis*. It differs from other populations of *castaneicollis* by having the throat and belly cream instead of white and reduced or absent chestnut colouration and U-patterning on the neck and flanks. Despite these genetic and morphological differences, *atrifrons* has similar vocalizations, habits and habitat to other forms of *P. castaneicollis* [2, 54], Hence, in terms of our stated criteria, its elevation to full species is not supported. The putative subspecies from Somalia, ‘*ogoensis*‘, is clinally more grey [2], and those from isolated populations west of Lake Zwai, ‘*kaffanus*’ are clinally less well-defined and U-patterned [2]. Moreover, their CYTB divergence from nominate *castaneicollis* is 0.2-0.7%. Hence, these taxa are synonymized within *castaneicollis*.

*Pternistis jacksoni* occurs between 2200 and 3700 m [54], primarily in forests, forest edges, moorlands, bamboo patches and within the Aberdares and Mt. Kenya, Mau Escarpment and Cherangani Mountains in *Podocarpus, Juniperus* and other Afro-alpine forests of western and central Kenya, extending marginally into Uganda (Fig 3). Jackson‘s Spurfowl is large (∼1130–1160 g, with females slightly smaller [54]). It has a red bill, yellow-orange ear-patch and eye-ring and tarsi with 1-2 spurs (upper shorter), only in the male. Its throat is buff and the lower neck greyish with the proximal part of the lower neck similarly patterned to the rest of the belly. Lower neck feathers are chestnut-coloured edged with buff to white, but the degree of chestnut and buff and white varies among individuals [2]. The subspecies ‘*pollenorum*’ from Mt. Kenya is not recognized because it is only somewhat darker [2] than other forms of *P. jacksoni*.

The closest CYTB taxon is *P. griseostriatus* at 5.0% sequence divergence.

### The Scaly spurfowls

The paraphyletic Scaly spurfowls comprise three allopatric species (*P. squamatus, P. ahantensis* and *P. griseostriatus*). The fourth species, *P. schuetti,* is parapatric with *P. squamatus* [Fig.4] [2]. Scaly spurfowls have the plainest plumage [2], with the least patterning and no strong colour. They are characterized by having ‘scaly’ underparts, and inhabit vestigial patches of montane and lowland forest, secondary and riverine forests, forest edges and clearing/cultivation therein of West Africa eastwards to the Sudan and north-eastern Tanzania, and Central Africa and the Benguela district of north-western southern Africa (Fig 4) [54].

**Fig 4.**
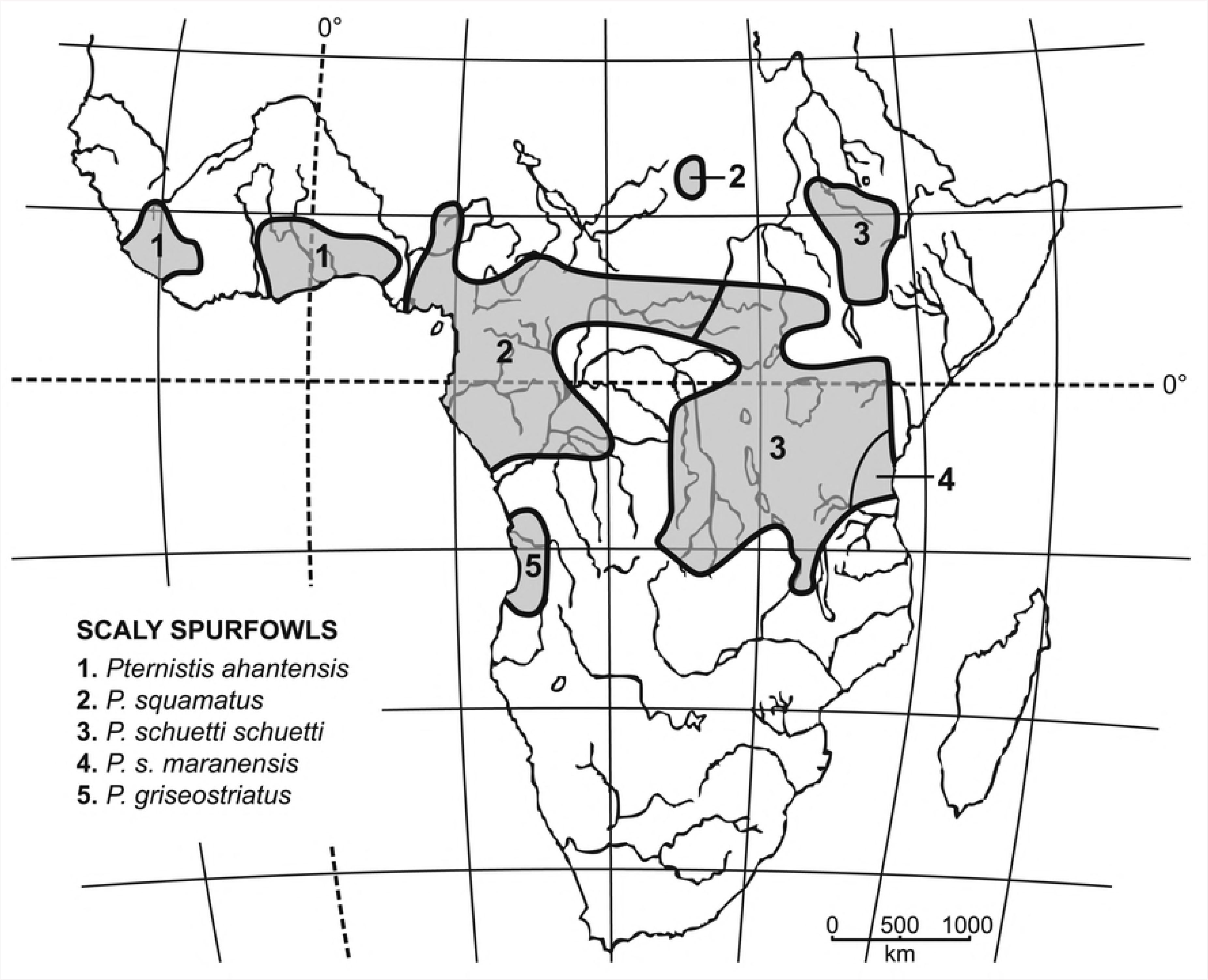
Geograpical distributions of Scaly spurfowls.

Compared with other spurfowls, these taxa are poorly diagnosed in terms of plumage pattern and colouration [54]. All taxa have unpatterned faces, whitish throats and brown upperparts, some with faint vermiculations. The underparts are brown or creamy-buff with very narrow darker edges, providing the characteristic ‘scaly’ appearance. There is no marked plumage dimorphism, with the exception that females tend to be more vermiculated than males [2].

*Pternistis squamatus* is sister to *P. schuetti,* and occurs in forested areas in south-eastern Nigeria, extending east into the DR Congo and up to 3000 m (on Mt Elgon) in Uganda/Kenya (Fig 4) [54]. It has a red bill, orange-red tarsi with 1-2 spurs (lower longer) in males only [54]. There is no size dimorphism (males 372-565 g., females 377-515 g.) and plumage, with U-patterned vermiculated upperparts, less so in males. It is the least distinctly patterned scaly taxon. The brown upperparts are indistinctly vermiculated faint grey with each feather with a blackish centre tinged maroon, and the upper back has faint buff U-patterning. The scaly underparts are brown with ill-defined dark shaft streaking [2].

The closest CYTB taxon is its sister-species, *P. schuetti,* at 3.4% sequence divergence.

*Pternistis schuetti* occurs in eastern DRC extending east to Uganda, Ethiopia, Kenya, Tanzania and Malawi [54] (Fig 4). It resembles *squamatus,* but is less vermiculated overall, and the scaly pattern on the lower neck is less clearly defined, each feather has a deep red-brown centre [2]. Populations west of the Rift Valley in Kenya south towards Kilimanjaro, Monduli and Mt. Meru in northeastern Tanzania [2], become clinally increasingly darker and greyer (more readily seen in males), and tend to have less white on the belly. Poorly sampled, isolated populations to the south ‘*usumbarae*‘, ‘*uzungwensis*’ and ‘*doni*’ are clinal or idiosyncratic variants of *schuetti*, but may warrant subspecific status should they exhibit significant genetic divergence.

The closest CYTB taxon to *schuetti* is *griseostriatus* at 2.7% sequence divergence.

*Pternistis s. maranensis* (1.2% divergent from nominate *schuetti*) occurs further east on Mt Kilimanjaro (up to 2000 m), Monduli, Mt Meru and in the Chyulu Hills (Fig 4). It is much darker and less patterned than *schuetti* [2]. There are scattered populations of Scaly spurfowls that show variation in plumage. About 240 km southeast of Kilimanjaro, birds (’*usambarae*‘) from the Usambara Mountains [2] have the areas around their eyes and cheeks freckled with black and white instead of uniform brown. Another isolated population from forests on the Vipya Plateau between 900 and 2800 m (’*doni*‘) in Malawi has upper and underparts that are more red-brown with some white streaking on the underparts [2]. These, for now, are included within nominate *schuetti*.

*Pternistis ahantensis* [54] occurs within gallery and secondary, coastal lowland West African forests in three disjunct populations west of the Niger River: southern Senegambia and northern Guinea-Bissau; southern Guinea, Sierre Leone and western Liberia; and north-eastern Ivory Coast and Ghana through the central Togo and central Benin to south-western Nigeria (Fig 4).

The Ahanta Spurfowl is a medium-sized spurfowl (males +-608 g., females +-487 g.) and has an orange bill with a black base and yellow-orange tarsi with 1-2 spurs (lower longer), only in the male [54]. It is the most patterned Scaly spurfowl, with breast and flank feathers having paler edges and darker centres. The feathers on its upperparts are vermiculated (distinct on the lower neck, indistinct on the back) with blackish centres and a reddish-brown shaft-streaking, those on the lower neck have some white U-patterning. The underparts are dark-brown chestnut with white and darker brown U-patterning [2]. The isolated western populations (’*hopkinsoni*’ for which we had no CYTB information) are paler overall [2] than those in the east and probably do not warrant taxonomic status.

The closest CYTB taxon to *ahantensis* is *P. squamatus* at 4.2% sequence divergence.

*Pternistis griseostriatus* is a small spurfowl (males 265-430 g., females 213-350 g. [54]) endemic to vestigial patches of forest in the Angolan western escarpment (Fig 4). The Grey-striped Spurfowl has a black bill with a red base (lower mandible orange-red) and its tarsi are orange-red with a single spur in the male. It is sexually monomorphic, and its lower neck feathers and wing coverts are chestnut and broadly edged and vermiculated with grey, similar to the pattern in *squamatus* and *ahantensis*, but paler. However, the underparts are plain, and the upper belly and flank feathers are chestnut and edged with greyish or creamy buff [2].

The closest CYTB taxon is *P. schuetti* at 2.7% sequence divergence.

### The Vermiculated spurfowls

Hall‘s Vermiculated taxa [2] are the most widely distributed spurfowls within Africa. They occur more or less continuously from Senegal to Eritrea southwards to Namibia and South Africa (Figs 5 and 6). There is even an isolated population (*ayesha*) of *bicalcaratus* in Morocco, making it one of the few sub-Saharan bird species with natural populations north and south of the Sahara [56]. Northern taxa frequent grasslands and cultivation within woodlands and acacia savanna and steppe. South of the equator, Vermiculated taxa frequent thick bush on hillsides and riparian watercourses. All taxa have brown or grey-brown heads, backs, wings and tails, with lighter vermiculations and/or V- and U-shaped patterning. Most taxa have a white eye-stripe.

**Fig 5.**
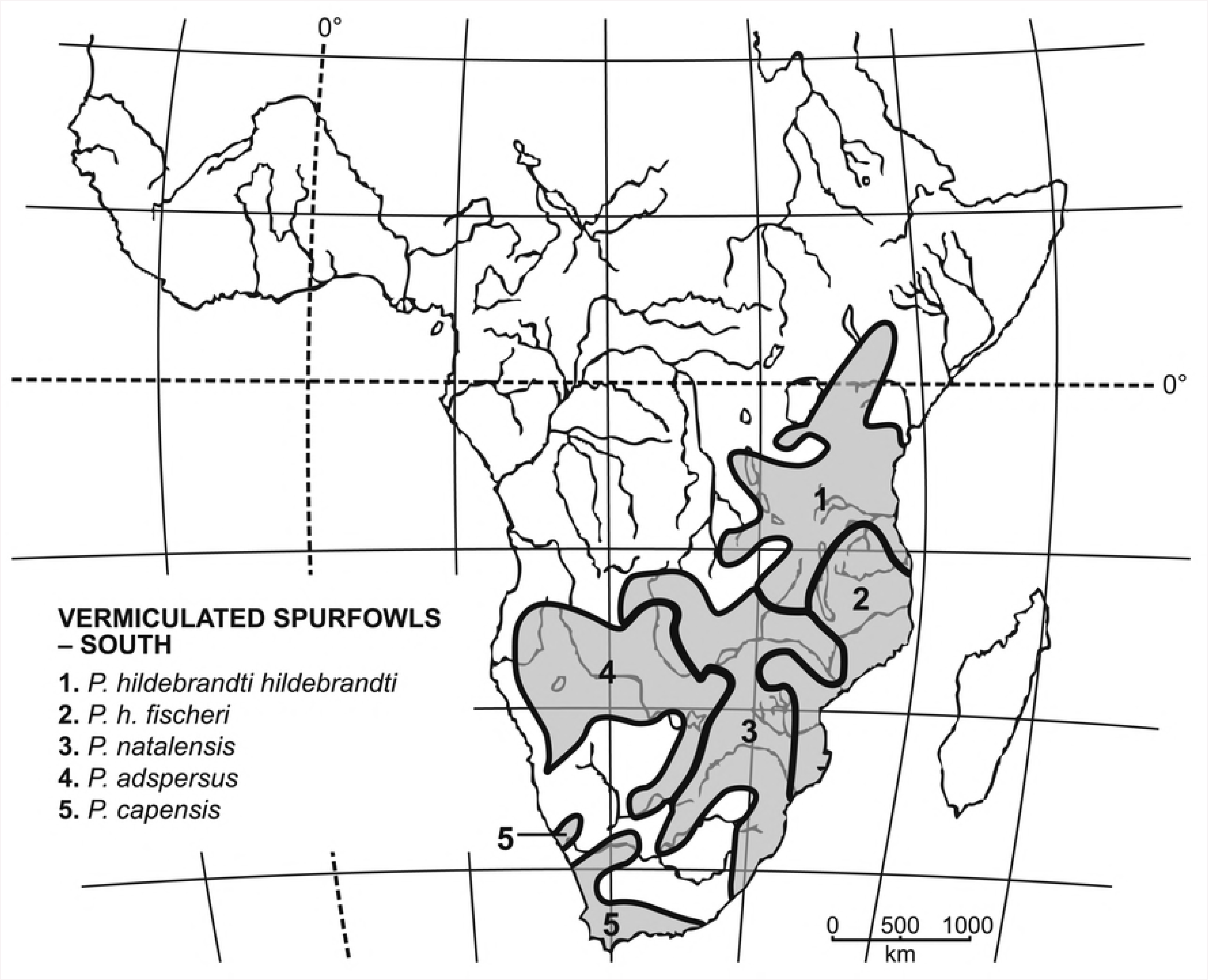
Geographical distributions of Vermiculated spurfowls (SOUTH).

**Fig 6.**
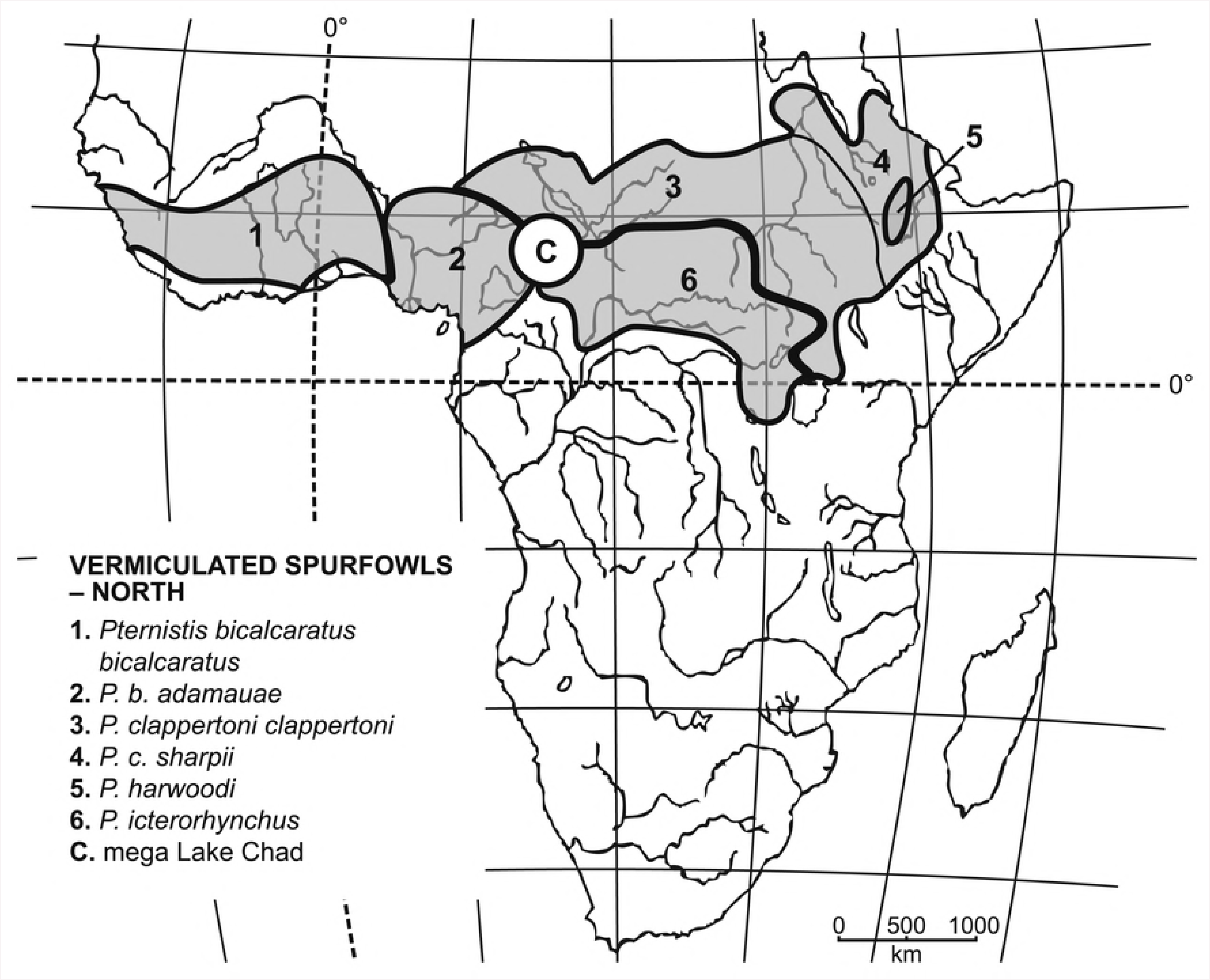
Geographical distributions of Vermiculated spurfowls (NORTH).

There are eight species and seven subspecies: *P. hildebrandti* (*hildebrandti* and *fischeri*), *P. natalensis, P. adspersus*; *P. capensis, P. icterorhynchus, P. bicalcaratus* (*bicalcaratus, adamauae, ayesha*), *P. clappertoni* (*clappertoni, sharpii*), and *P. harwoodi*.

Within the southern taxa, *P. hildebrandti,* occurs from sea level to about 2500 m. in east and south-central Africa, east and south from Lake Victoria through Kenya, most of Tanzania, northern Mozambique, north-eastern Zambia and Malawi (Fig 5) [54]. The species is sparsely distributed on rocky ground associated with dense thicket along rivers and on hillsides, acacia savanna, Miombo woodland and forest edge.

Hildebrandt‘s Spurfowl is a medium-sized spurfowl (two males 600 and 645 g., two females 430 and 480 g. [54]) and comprises two subspecies (*hildebrandti* and *fischeri*) with the former being sexually dimorphic. It has a reddish mandible and brown culmen with a yellow base, a yellow ear-patch and eye-ring, red tarsi with 1-2 spurs on both sexes. The dorsal plumage of males resembles that of *P. icterorhynchus*. It is greyish brown with vermiculations, and the hind and lower neck are streaked black with white margins, and the belly plumage has marked black blotching. Females have similar back plumage to males, but (especially in *P. h. fischeri*) differ markedly in having orange-brown underparts [2].

*Pternistis h. fischeri* [2] (1.0% sequence divergent from *hildebrandti*) from southern Malawi, Mozambique and south-western Tanzania differs from *hildebrandti* in that females have an unpatterned nape, hind neck and upper belly, in sharp contrast to an orange-brown abdomen. Birds from Kenya, ‘*altumi*’ [2], do not warrant taxonomic recognition because their plumage is intermediate between nominate *hildebrandti* and *fischeri*.

The sister-taxon to *P. hildebrandti* is *P. h. fischeri* with a CYTB (for *fischeri*) sequence divergence of 0.8%. Other forms of *hildebrandti* are >2% divergent from *natalensis*, its sister-species. In the Luangwa valley, the presence of specimens with intermediate plumage suggests that *P. hildebrandti* may interbreed (or have interbred) with *P. natalensis* [2].

*Pternistis natalensis* is a medium-sized spurfowl (males 415-723 g. females 370-482 g. [54]) distributed across south-eastern Africa, from Zambia, Zimbabwe, inland Mozambique, eastern Botswana, Swaziland and north-eastern South Africa (Fig 5). It occurs in thick riverine bush, but will venture into dry lowveld savanna and adjacent grasslands [9]. The Natal Spurfowl [54] has an orange bill with a dull greenish base, and the orange tarsi have a single spur, only in the male. It is normally sexually monomorphic, but some populations ‘*neavei*’ from southern Zambia and western Mozambique are slightly dimorphic. The hindneck is mottled black and white, the back is highly vermiculated in greyish-brown and black, with white and buff markings. The belly is buff with the upper belly to mid-belly being heavily patterned in black and buff U-patterning is concentrated on the breast with the extreme lower abdomen having no or few marks [2].

The closest CYTB taxon is *P. hildebrandti fischeri* at 0.8% sequence divergence. Next closest is *P. h. hildebrandti* at 3.1% divergence.

*Pternistis adspersus* is a smallish spurfowl (males 340-635 g., females 340-549 g. [9, 54]) and occurs in dense bush, mixed woodland and low scrub thickets interspersed with open ground, mostly on Kalahari sands along watercourses in Namibia, Botswana, southern Angola and south-western Zambia (Fig 5). The Red-billed Spurfowl is a monotypic species with an orange-red bill and tarsi, yellow ear-patch and eye-ring. Males have a single spur. The upperparts are finely vermiculated, and the underparts are narrowly distinctly barred with black and white, variably on the lower neck [2].

The closest CYTB taxon is *P. capensis* at 3.8% sequence divergence.

The monotypic *Pternistis capensis* is the largest Vermiculated spurfowl (males 870-1000 g., females 640-900 g.). It is endemic to thick cover and rocky river valleys in the Fynbos Biome of south-western South Africa, with isolated populations extending deep into the Karoo biome and lower stretches of the Orange River (Fig 5) [9]. The Cape Spurfowl [9] has a brown upper mandible (lower red), and orange red tarsi with one spur (females) and sometimes two (males). It has distinctive uniform brown and white double V- or U-shaped patterning on the back, breast and belly, while the throat has irregular black flecking. The breast and belly feathers have broad white shaft streaks [2].

The closest CYTB taxon is *P. adspersus* at 3.8% sequence divergence.

Moving to the northern vermiculated taxa, *P. icterorhynchus* is a medium-sized spurfowl (males 504-588 g. females 20-462 g. [54]) and occurs in grasslands, open woodlands and adjacent agricultural lands in the Central African Republic, northern DR Congo, extending east to South Sudan and Uganda (Fig 6). Heuglin‘s Spurfowl has a yellow-orange black bill, small yellow eye-patch, yellow-orange tarsi with 1-2 (upper longer), in males only. It is monotypic and sexually monomorphic species (Fig 6), with a chestnut crown, brown back diagnosed by having less V-shaped patterning on the lower neck and more vermiculations on the back than other vermiculated taxa. Its underparts are buff heavily marked with dark brownish-back V-shaped markings [2].

The closest CYTB taxon is *P. bicalcaratus* at 3.3% sequence divergence.

*Pternistis bicalcaratus* comprises three sexually monomorphic subspecies (Fig 6). All the taxa are similarly patterned above and below, differing in the degree of colouration and vermiculation, and the size of the arrow-shaped buff marks in the centre of the belly feathers [2]. They occur [54] in dry grasslands, open savanna, palm groves and cultivated areas of West Africa from Senegal east to northern Cameroon and southern Chad (Fig 6). The nominate form of the Double-spurred Spurfowl [54], *bicalcaratus* is a medium-sized (males +-507 g., females +-381 g.) spurfowl, and has a greenish-black bill and 1-2 greenish tarsi (upper longer), much shorter in females. It has no bare facial skin. It has a pale rufous crown, and a white eyestripe. It has rufous-chestnut on the lower neck and the remaining upperparts are vermiculated with V-shaped patterning. It has buff underparts, distinctly and heavily streaked with black and chestnut small arrow-shaped buff marks on most belly feathers [2]. The more heavily patterned *ayesha* (from Morocco, not mapped) is similar (1.0% CYTB divergent) to *bicalcaratus*, but is faintly vermiculated and slightly more rufous on the lower neck, with small arrow-shaped buff marks on the belly feathers [2]. The darkest form is *adamauae* (1.7% CYTB divergent) with very little rufous on the lower neck, and the underparts are more buff with extremely reduced chestnut and larger arrow-shaped buff marks long the centres of the belly feathers [2].

The closest CYTB species is its sister-species, *P. icterorhynchus,* at 2.7% sequence divergence.

*Pternistis clappertoni* [54] comprises two widespread subspecies extending up to 2300 m in semi-arid grassland and bushy savanna and adjacent cultivations across north-central Africa from far eastern Mali, central Niger, far north-eastern Nigeria, Chad, southern Sudan, South Sudan, northeastern Uganda and western Ethiopia (Fig 6). It also occurs in the Nile and Blue Nile River valleys [2].

Clapperton‘s Spurfowl is a medium-sized spurfowl (males 450-604 g. females 300-530 g. [54]). It has a black bill with a red base and red tarsi with 1-2 spurs in males only. The bare skin around the eye distinguish it from *P. bicalcaratus, P. icterorhynchus* and *P. castaneicollis*. The brown upperparts of the nominate form, *clappertoni*, have U-shaped patterning (very similar to those of *P. icterorhynchus*), but are more orange brown and vary geographically in the degree of vermiculation and U-patterning. It has a fairly extensive white throat and the neck is buff below with black to brownish marks. *P. c. sharpii* (1.4% CYTB divergent from *clappertoni*) has marks on the belly which are streakier than those in *clappertoni* in having a more buffy white background below with the upper belly being similarly U-patterned extending onto the back [2]. A single specimen collected at “Ngeem” at Lake Chad (possibly Nguigmi), the type of *Francolinus’ tschadensis‘* is possibly a hybrid between *clappertoni* and *icterorhynchus* [2].

The closest CYTB taxon to *clappertoni* is its sister-speces, *P. harwoodi,* at 1.4% sequence divergence. The next closest taxon is *P. bicalcaratus,* jumping to 3.1% divergence.

*Pternistis harwoodi* is a medium-sized (one male 545 g., one female 446 g. [54]), poorly known species occurring in *Tyhpa* reedbeds, scrub, thicket and adjacent cultivations along the gorges of the Jemmu valley, the Blue Nile and its tributaries of East Africa, and the highlands of central Ethiopia (Fig 6).

Harwood‘s Spurfowl [2] has a red bill with a black tip, bare red eye-ring and tarsi with 1-2 spurs, in males only. It most closely resembles *P. natalensis*, which lacks the bare red facial skin, but has more defined U-patterning on the nape, with similar U-patterning on the underparts. The upperparts of the male that we examined is grey speckled and finely barred with blackish and buff above. The lack of a white eyestripe sets it apart from other Vermiculated spurfowls. The hind and lower neck, sides of face, and throat are speckled with black and white. It has irregular double-V shaped patterning on its underparts which tends to be scattered on the lower extreme of the buff belly.

The closest CYTB taxon is *P. clappertoni* ‘*sharpii*’ at 0.7% sequence divergence. The next closest taxon is *P. natalensis* at 4.8%.

### The Bare-throated spurfowls

We recognize five species and five subspecies: *leucoscepus* (*leucoscepus, infuscatus*), *cranchii, afer* (*afer, castaneiventer, humboldtii*), *swainsonii,* and *rufopictus.*

The Bare-throated spurfowls [2, 54] are largely allo/parapatric and ecologically segregated meta-populations, extending from Ethiopia and Eritrea in northeast Africa, westwards through Kenya, Tanzania, Sudan and Uganda to the Congo and Gabon, and south through Angola, northern Namibia, Botswana, Zimbabwe, and Mozambique to South Africa (Figs 7 and 8). Species inhabit mesic lowland grasslands and open woodland savanna/bush often adjacent to water.

**Fig 7.**
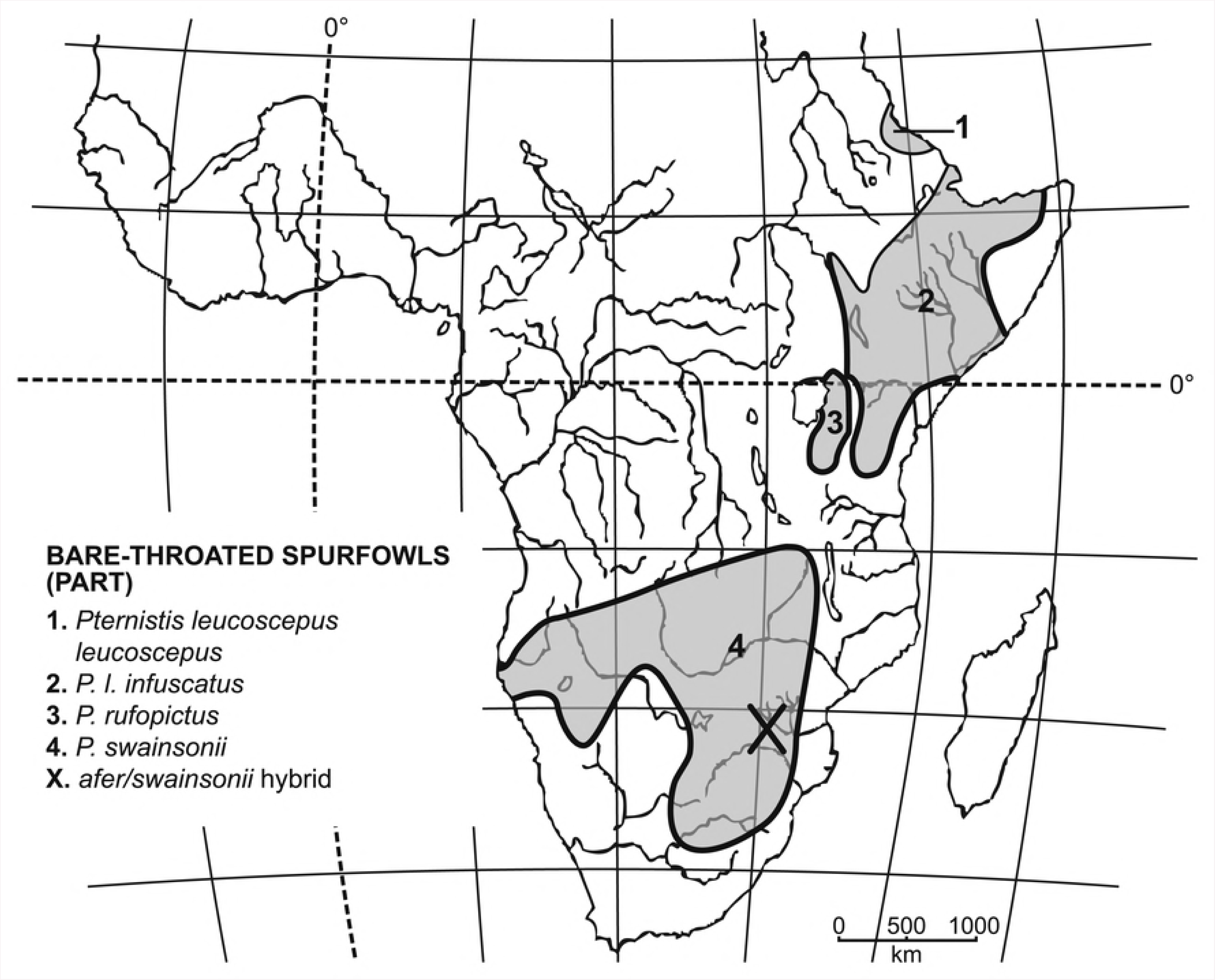
Geographical distributions of Bare-throated spurfowls (part).

**Fig 8.**
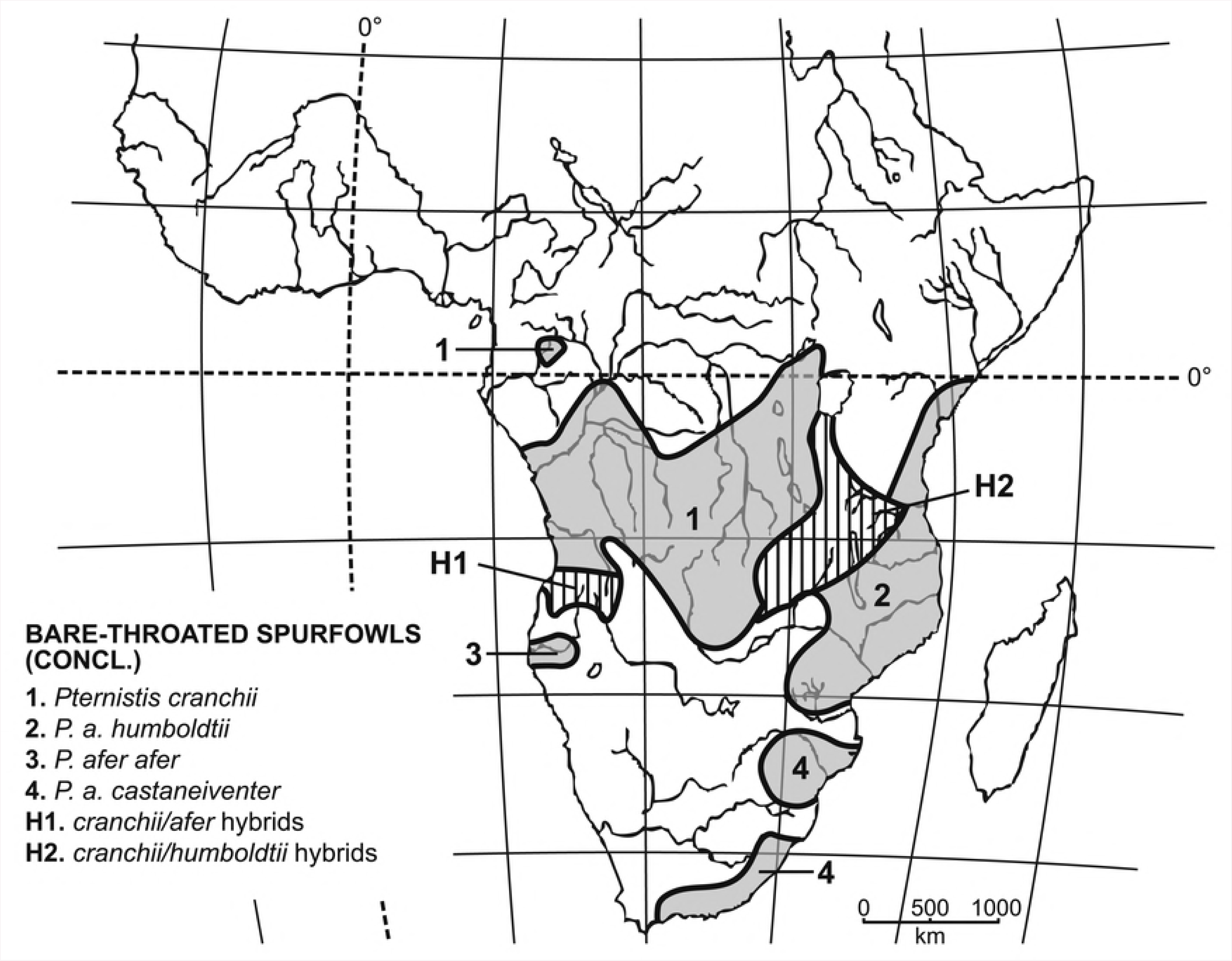
Geographical distributions of Bare-throated spurfowls (concl.).

Bare-throated spurfowls are sexually monomorphic in plumage (although females of some species are slightly vermiculated), with a body mass ranging from 340–950 g) [2, 54]. They are distinguished from other spurfowls by having bare skin on the throat and a patch around the eye and plain dark upperparts without pale vermiculations. Their tarsi are black, red, orange or brown with spurs well-developed in males only. They have a long robust lower spur and, in some taxa (*P. leucoscepus* and *P. rufopictus*), often a shorter blunt upper spur, less prevalent in *P. afer* and rare in *P. swainsonii* [2]. *Pternistis leucoscepus* is a medium-sized, markedly dimorphic spurfowl (males 615-896 g., females 400-615 g. [54]). This species is the most morphologically and ecologically differentiated species and comprises two subspecies: *leucoscepus* and *infuscatus* (Fig 3). It occurs in arid-acacia savanna and sub-desert scrub in eastern Africa (most of Kenya, north-eastern Uganda, south-eastern South Sudan and northern Tanzania), extending north and east through Ethiopia and Somalia nearly to the tip of the ‘Horn’ of Africa (Fig 7) [2, 54].

Both subspecies of the Yellow-necked Spurfowl [2, 54] have black bills with a red base, bare red skin around the eye, bare yellow throat skin, and black tarsi with 1-2 spurs on the males. The upper back plumage is dark brown with white shaft streaks and the underparts are streaked with white and chestnut with narrow white edges and a triangular white patch at the tip, tapering up the shaft. The primaries have a conspicuous white patch, which is visible during flight. The northern subspecies *infuscatus* at 0.9% sequence divergence from *P. l. leucoscepus*, differs in having more chestnut than white on the underparts in contrast with the dominant white over chestnut present in *leucoscepus*.

The closest CYTB taxon to *P. leucoscepus* is *P. cranchii* at 3.5% sequence divergence.

Hall‘s [5] ‘Red-necked’ Spurfowl [2, 54] is the most widespread and morphologically variable ‘species’ of Bare-throated spurfowl. It has a complex geographical distribution and occurs in relatively mesic evergreen forest edges, and woodland in central Africa and Kenya, extending southwards through, Zambia, Malawi, Tanzania, south-western Angola, north-western Namibia, eastern Zimbabwe, Mozambique into eastern South Africa (Fig 8) [9].

All taxa ascribed to this spurfowl were lumped into one species [2], *afer*, with two polytypic subspecies, *afer* and *cranchii* [h]. We elevate *cranchii* to full species status (Fig. 8). Both species are medium-sized (males 480-1000 g., females 370-690 g.) and have a red bill, throat skin and tarsi with 1-2 spurs in males only [9].

*Pternistis cranchii* [2, 9] includes populations from southern Congo, northern Angola, northern Zambia, western Tanzania, Uganda and Lake Victoria shores (Fig 8). It is characterized by having no white on the head or black on the abdomen. The underparts are heavily and finely vermiculated with grey with heavy chestnut brown streaking on the abdomen. Its lower belly feathers have buff central streaks vermiculated with blackish grey and margined with broad chestnut (degree of chestnut colour varies geographically) and a black and grey facial pattern. Populations from the Ruzizi valley, north of Lake Tanganyika, (’*harterti*‘) are much darker overall, and the streaking on the abdomen is maroon, rather than chestnut [2].

Within Hall‘s [2] ‘cranchii’ forms (*cranchii*, ‘*nyanzae*‘, ‘*harterti*‘), the CYTB divergences are c. 1%, and the lowest CYTB divergence between a form of *cranchii sensu lato* to one within *afer* (from Angola) is 1.6% sequence divergence. Thereafter, the pairwise divergence values for forms of *cranchii* versus *afer* well exceed 2%. All ‘hybrid’ forms studied (’*intercedens*‘, ‘*bohmi*‘, ‘*itigi*‘, ‘*cunenensis*’ and ‘*benguellensis*‘) are 0.7-0.8 % divergent from ‘pure’ *cranchii* and >2% divergent from *afer* taxa.

In marked contrast to *cranchii*, populations of *P. afer sensu stricto* [2, 9, 54] have unvermiculated underparts, and are strongly patterned black and white on the face and underparts, which have broadly streaked greyish black central streaks with buff margins (particularly in the nominate subspecies *afer*) or have thin greyish black central streaks separating the long buff parallel streaks margined with black or sometimes maroon (in south-eastern South African specimens).

In the nominate *P. a. afer,* confined to south-western Angola (Fig 8), the face is white, and the underparts are streaked broadly with black and white, with black centres and white margins. Elsewhere in Angola, specimens from the Upper Okavango basin generally resemble *cranchii* with some intermediate plumage forms ‘hybrids’ – ‘*cunenensis*’ genetically closest (0.5% divergent) to *cranchii*.

The closest CYTB species to Angolan *P. a. afer* is *P.* c. *cranchii* at 1.6% sequence divergence. It is 2.7% divergent from South African *P. afer*.

In south-eastern Africa, *P. a. castaneiventer*, occurs in South Africa from KwaZulu-Natal south and west into the Eastern Cape Province (Fig. 8). It has a wholly black face with the upper breast and abdomen streaked with black and white, edged maroon in birds from KwaZulu-Natal. Birds from eastern Zimbabwe and southern Mozambique have a white face and ‘necklace’ above the breast.

*P. a. castaneiventer* is 2.7% divergent from South African *P. afer*.

The closest non-*afer* CYTB species-level taxon is the *P*. *c. ‘cunenensis‘* at 2.7% sequence divergence.

*Pternistis a. humboldtii* ranges within eastern Africa, from southern Kenya and Tanzania south to Mozambique (Fig. 8). It is 1.3% divergent from *P. a. castaneiventer* and 2.6% from *P. cranchii*, has a black face with a white jaw-beard and black belly patch. Feathers on the upper belly are mainly grey with black shaft streaks which contrast with the abdomen to form a black patch, and the flanks which are streaked black and white. Birds from coastal Kenya, have a white face and black and white eyestripe. Birds from northern Tanzania southwards to Malawi and southeastern Zambia have a wholly black face [2].

A range of additional idiosyncratic subspecies of intermediate phenotype relative to the *cranchii* and *afer* have been described where these three forms are para/sympatric, but they lack the morphological cohesion necessary for recognition. These occur mainly in two hybrid zones between *cranchii* and *afer*. One stretches from Kondoa Dodoma in central Tanzania through central Malawi into the Luangwa Valley (Fig. 8). Hybrids have well-defined streaks on the abdomen and varying amounts of chestnut and black-and-white depending on relative proximity to the respective parental forms, but show little within-locality morphological variation [2]. The second hybrid zone in northern and central Angola is characterized by morphologically relatively unstructured populations [2].

*Pternistis rufopictus* is a monotypic medium-large spurfowl (males 779-964 g., females 400-666 g. [2, 54]) distributed in dry savannas, thickets and plains from the south-eastern shores of Lake Victoria to the Wembere River in north-western Tanzania (Fig 7). It is narrowly sympatric with *P. leucoscepus* where their distribution overlaps in the southern parts of its range [2].

It has a red bill, orange-pink throat skin, bare red skin around the eye, and brown tarsi with 1-2 spurs on males only. The eye-stripe and sides of the face are black and white. It also has a white chin stripe on either side of the bare throat. Its upper back plumage is grey-brown with dark vermiculations and dark shaft streaks, grading posteriorly to black, white and chestnut streaking. The wing coverts and feathers on the back are edged with rufous chestnut. The upper belly is grey with black shaft streaks and the lower belly is streaked black and white. The lower belly feathers have narrow central black streaks separated from rufous chestnut margins by broad buff to white streaks. *P. rufopictus* is similar to the *cranchii*-type taxa in western Tanzania, except for the white chin stripes on either side of the throat (as in *humboldtii*), and no vermiculations [2].

Its closest CYTB taxon is *P. afer cranchii* at 1.7% sequence divergence. Its next closest CYTB taxon is *P. leucoscepus* at 4.0%.

*Pternistis swainsonii*, is a monotypic, medium-sized spurfowl (males 400-875 g., females 340-750 [9]) distributed across south-western Africa from northern Namibia, eastern Botswana, Zimbabwe, southern and eastern Zambia, southwards to north-eastern South Africa (Fig 7). It frequents acacia/mopane savanna and tall grassland, almost anywhere where there is suitable cover. It is especially partial to cultivated lands. Its range and numbers have increased in recent decades in the south-eastern parts of its distribution due to agriculture-related alteration of the environment.

Swainson‘s Spurfowl [9] has a black upper mandible (lower dull orange), bare red throat skin and black tarsi, normally with a single spur in the male. Its upperparts are grey brown with faint dark shaft-streaking. The underparts are similar but with a grey wash on the breast and chestnut streaking lower down. Specimens from southern Zimbabwe and northern South Africa have blackish mottling on the abdomen. The feathers have a narrow central greyish black streak separated from greyish chestnut margins by broad buff grey vermiculated streaks [2].

The closest CYTB taxon to *P. swainsonii* is *P. cranchii* at 3.6% sequence divergence.

## Discussion

### Origin of African spurfowls and ‘groups’

The African spurfowls represent a remarkable biogeographical, morphological, behavioural and ecological radiation within the entire African continent. The existence of a subspecies of *Pternistis bicalcaratus* in Morocco, exceptional amongst Afrotropical birds [56], demonstrates relatively recent biogeographic connectivity between North and sub-Saharan Africa.

African spurfowls are sister to *Ammoperdix heyi* [native range from Egypt and Israel east to southern Arabia] and *Perdicula asiatica* [native range India, Nepal, Bangladesh, Pakistan and Sri Lanka] [20], which are both arid-zone taxa [8]. The phylogenetically most basal African spurfowl, *hartlaubi*, is also a highly peculiar, desertic bird [9, 53]. Therefore, African spurfowls may have been derived from an arid-adapted taxon that dispersed from the Middle East or Asia into Africa (30-40 mybp, 20] during a continent-wide arid era. Hall [2] also suggested an Asiatic origin. Within Africa, dispersal to the south may have been facilitated by an ‘Arid Corridor’ that has multiply connected the northeast arid Horn of Africa to arid Namibia and the Karoo in the southwest [57, 58, 59, 60].

### Montane and scaly spurfowls

The Montane and Scaly spurfowls follow on from *hartlaubi* paraphyletically (Fig 2). They are probably results of invasions of, and diversification within, forested biotopes where they predominated thereafter during subsequent wetter eras. Initially, when forests subsequently contracted geographically during renewed dry eras, proto-Montane spurfowls became isolated in relictual, island-like patches of montane forest. This scenario is supported by two of the relatively basal, most isolated, western Montane taxa (*camerunensis* and *swierstrai*) being geographically most proximal to the hill/mountain-dwelling *hartlaubi*, and relatively small, sexually dimorphic, and poorly spurred.

The Noble Spurfowl, *P. nobilis*, this Montane taxon is geographically intermediate between western and north-eastern African Montane taxa and is sister to *camerunensis* and phylogenetically ‘links’ all of these western taxa to those in the northeast. It is also of intermediate body mass between the two species assemblages [54].

The divergence of Scaly spurfowls from Montane taxa is probably more a consequence of ecologically opportunistic speciation during multiple expansions and contractions of lowland forests separated by intervening savanna/steppe, hence the relatively close genetic propinquity between montane *jacksoni* and scaly *griseostriatus*, and montane *swierstrai* and scaly *squamatus*.

Although the core ranges of *ahantensis* and *squamatus* closely coincide with the current distribution of present-day lowland forest, the existence of peripheral, island-like isolates suggests a much broader continuous distribution in more widespread forest during wetter eras. Indeed, the primordial ‘scaly’ spurfowl may have been a single species distributed continuously from West Africa eastwards to the East African coast and south to Angola, with an initial vicariance event producing *griseostriatus.* The second major forest vicariance event and physical barrier of the Niger River may have split *ahantensis* from *squamatus*.

Furthermore, vicariant ‘subspeciation’ within proto-*squamatus*, may have promoted the divergence of *schuetti* in paleo-forest isolates in the east (Fig 4) during drier eras, as it seems to have done within Latham‘s Forest Francolin, *Afrocolinus lathami* [Mandiwana-Neudani et al., in review] and Plumed Guineafowl, *Guttera plumifera* [61].

Finally, Hall [2] noted that *squamatus* extends its range to higher altitudes on mountains uninhabited by montane spurfowls, suggesting that competition might also limit its range.

### Vermiculated and Bare-throated Spurfowls

Moving into relatively open arid-steppe, savanna, woodland and bush biotopes, the vicariant speciation of Vermiculated and Bare-throated taxa was within pockets of these biotopes promoted by physical barriers (lakes, rivers and valleys), other geomorphological events and expansion and contraction of forest [62, 63].

For example, the southern Vermiculated *P. hildebrandti* and *natalensis* have similar habitats and are separated by the valleys of the Shire and Luangwa Rivers [2]. Within the northern Vermiculated taxa, Lake Chad probably played a similar role in speciation between proto-*bicalcaratus* and proto-*icterorhynchus* + *clappertoni* [62]. These latter two spurfowls perhaps diverged in broad stretches of arid (*clappertoni*) and mesic (*icterorhynchus*) savanna/grassland. Riverine forest along the Nile and in Kenya/Uganda could also have separated proto-*icterorhynchus* in the north from proto-*hildebrandti* in the south [2].

The initial divergence of Vermiculated taxa probably occurred in central/southern Africa with the proto-southern taxa radiating within the region into xeric western (*adspersus* + *capensis*) and mesic eastern (*hildebrandti* + *natalensis*) clades. Northern taxa may be a result of invasion from the south via the ‘Arid Corridor‘.

In sharp contrast, the northward dispersal of *bicalcaratus* from West Africa into Morocco was via a relatively recent corridor of savanna biotope that subsequently reverted to the western Sahara.

With regard to Bare-throated taxa, proto-*leucoscepus* originated in arid biotopes in the north and subsequently dispersed southwards, once again via the ‘Arid corridor‘, with proto-*cranchii*/*afer*/*swainsonii* biogeographically insinuating themselves within southern Vermiculated taxa.

Ecological speciation due to competition may also have contributed to speciation in Vermiculated taxa. Those north of the equator are birds of grasslands and cultivations in woodlands, savannas and steppe. But, south of the equator, these habitats are occupied by Bare-throated taxa, and southern Vermiculated taxa are relegated to thickets on rocky hillsides and along rivers.

### Relevance of the ‘Realm’ of Tokogeny

There is also evidence that tokogenetic processes may have played significant roles in the evolution of *Pternistis* species and subspecies. Interbreeding is most apparent in the Vermiculated and Bare-throated taxa which may continue to ‘hybridize’ in captivity or where they come into contact in nature. For example, where *P. cranchii* and *afer* hybridize along the ‘Arid corridor’ and especially in eastern Zambia west of the Luangwa River, south to 13°30' and also in the Eastern Province plateau in Lundazi, hybrid forms show remarkably high within-locality morphological homogeneity, forming microgeographic ‘races‘. But, where *cranchii* and *afer* hybridize in southern Angola and northern Namibia, there is no such morphological homogeneity. Indeed, Roberts [64] described a ‘new species’ of spurfowl, *P. cooperi,* from near Harare, Zimbabwe (Fig. 7), which turned out to be a hybrid between *cranchii* and *swainsonii*, probably due to range expansion by *swainsonii* into *cranchii* habitat which was transformed by agriculture.

McCarthy [29] also reports a range of spurfowl hybrids, mainly within and between Vermiculated and Bare-throated taxa: *afer* X *leucoscepus*; *afer* X *swainsonii*; *bicalcaratus* X *erckelii*; *castaneicollis* X *erckelii*; *hildebrandti* X *natalensis*; *leucoscepus* X *rufopictus*; *natalensis* X *swainsonii; adspersus* X *natalensis*, and *adspersus* X *swainsonii* recorded by Little [65, 66].

Perhaps the most interesting taxon in this regard is *P. rufopictus*, which Hall [2] speculated might have resulted from stabilized hybridization. This is because it is ‘diagnosed’ by a combination of characters of the other Bare-throated taxa (e.g. orange, rather than red or yellow facial skin) and ‘hybrid’ (vermiculated, chestnut, white and black) plumage. Genetically, it is +-4% divergent from *leucoscepus*, +-1.8% from cranchii and hybrids, and 2.4-3.2% from *afer.* Phylogenetically, it ‘links’ *swainsonii* + *cranchii* with *afer*. Vocally, it sounds very similar to *P. leucoscepus* except that its call is much ‘faster‘. The strophes of *P. leucoscepus* and *P. rufopictus* are both high-pitched, with an element of screeching and more protracted trilling [17]. Nevertheless, its specific status seems appropriate since it seems to exist partially sympatric with *leucoscepus* and *afer* without unfettered hybridization. Its putative hybrid origins remain to be tested using genomic data.

## Acknowledgements

We acknowledge ‘Pat’ Hall‘s contributions as a pioneering systematist/biogeographer and as a generous mentor. We thank the curators and collection managers at the American Museum of Natural History (New York), the Natural History Museum (Tring), Humboldt-Museum (Berlin), the National Museum of Natural History - Smithsonian Institution (Washington, D.C.), the Ditsong National Museum of Natural History (Ditsong Museums of South Africa), the Iziko Museum of Natural History, the Natural History Museum of Zimbabwe (Bulawayo), the Field Museum of Natural History (Chicago), Carnegie Museum of Natural History (Pittsburgh), Academy of Natural Sciences of Philadelphia, Natural History Museum of Los Angeles County, Museum of Comparative Zoology (Harvard University), and the Royal Museum for Central Africa (Turvuren) for access to specimens under their care. We thank Robert Moyle for help with designing primers and Julie Feinstein for helping with the sequencing of toe pads sub-sampled from museum skins. This project was funded by the South African Department of Science and Technology (DST) and the National Research Foundation (NRF) through their South African Biosystematics Initiative and Centre of Excellence Programmes, and the African Gamebird Research, Education and Development Trust. Amy Bruce kindly assisted with constructing the figures.

**Appendix 1.**
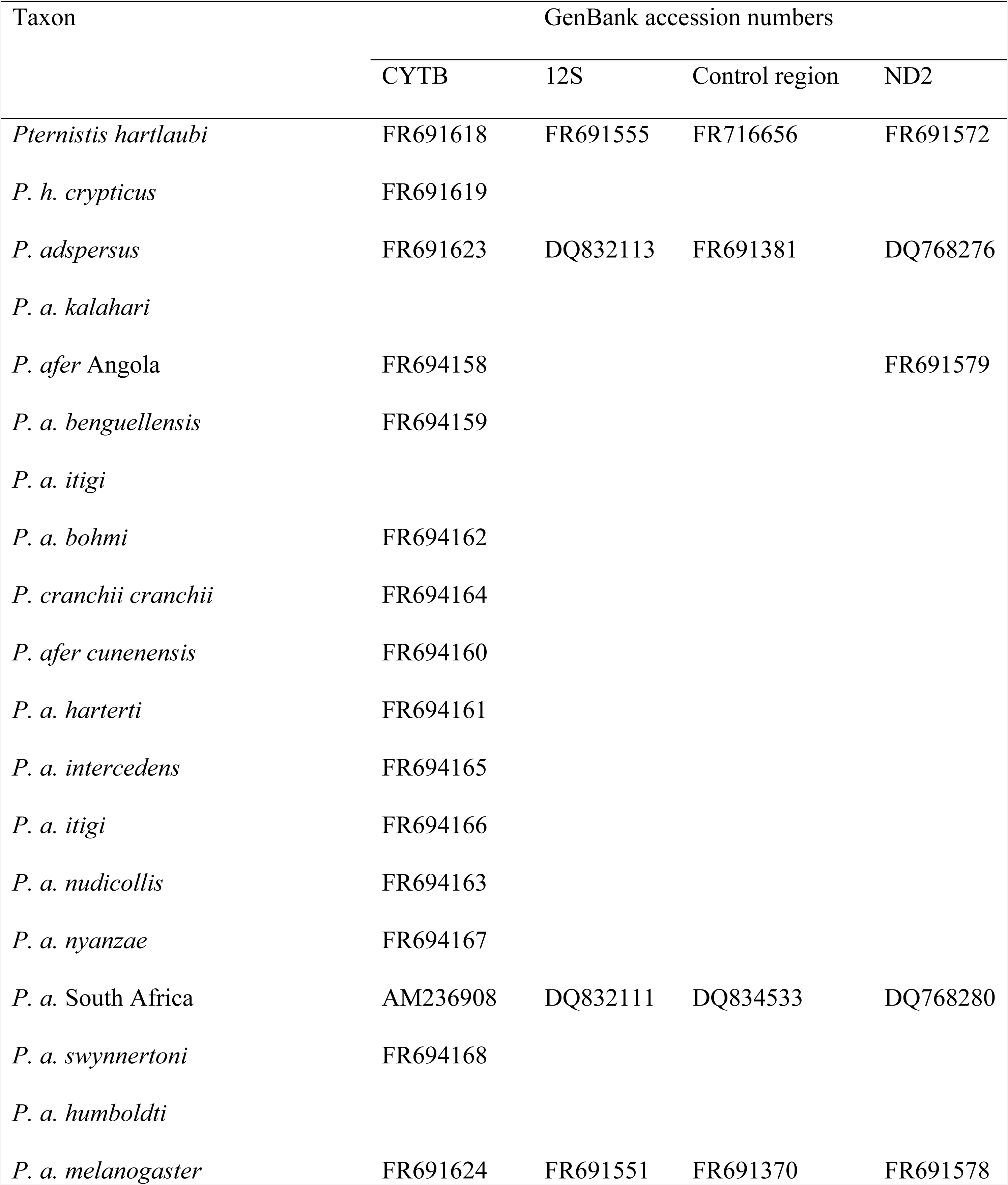
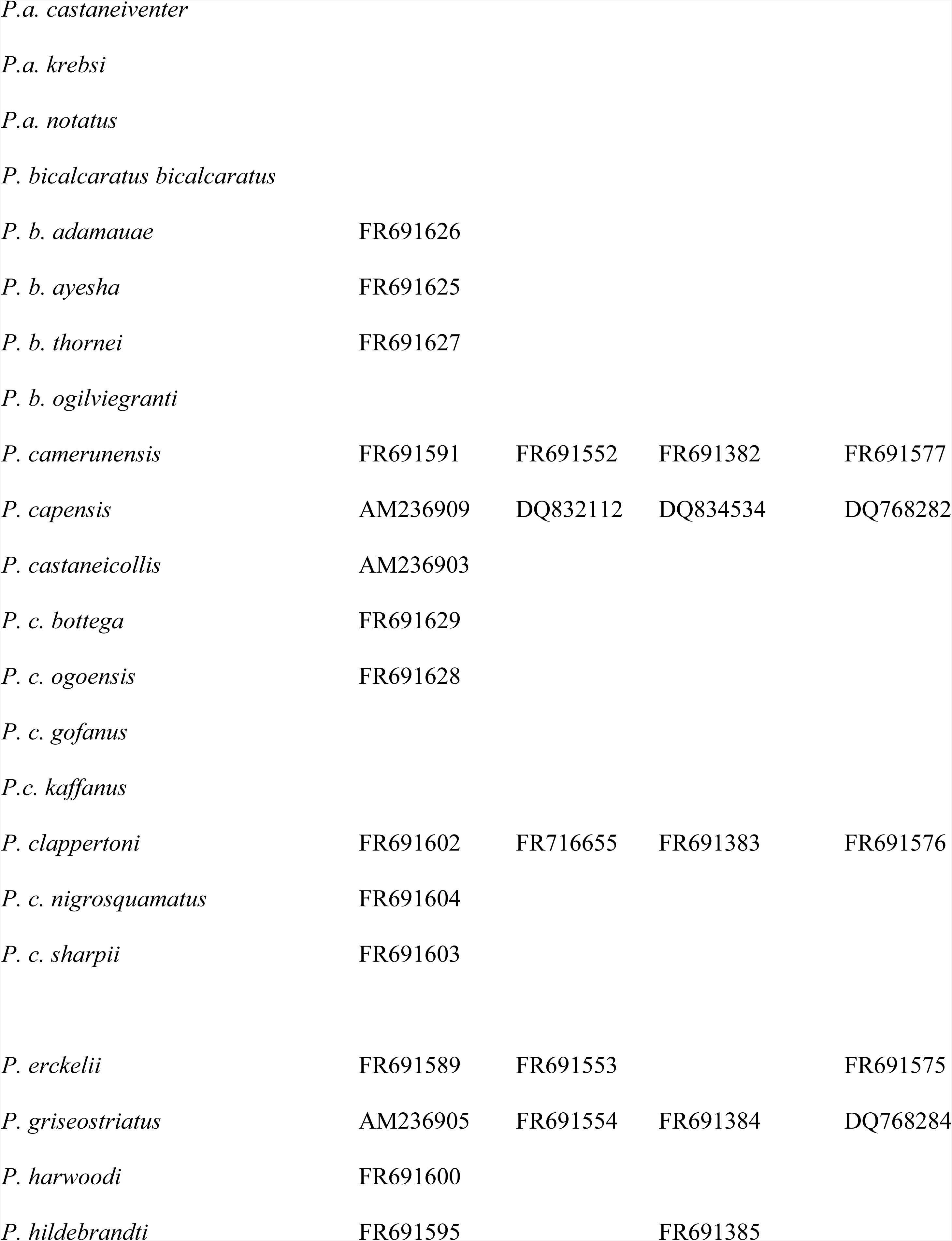
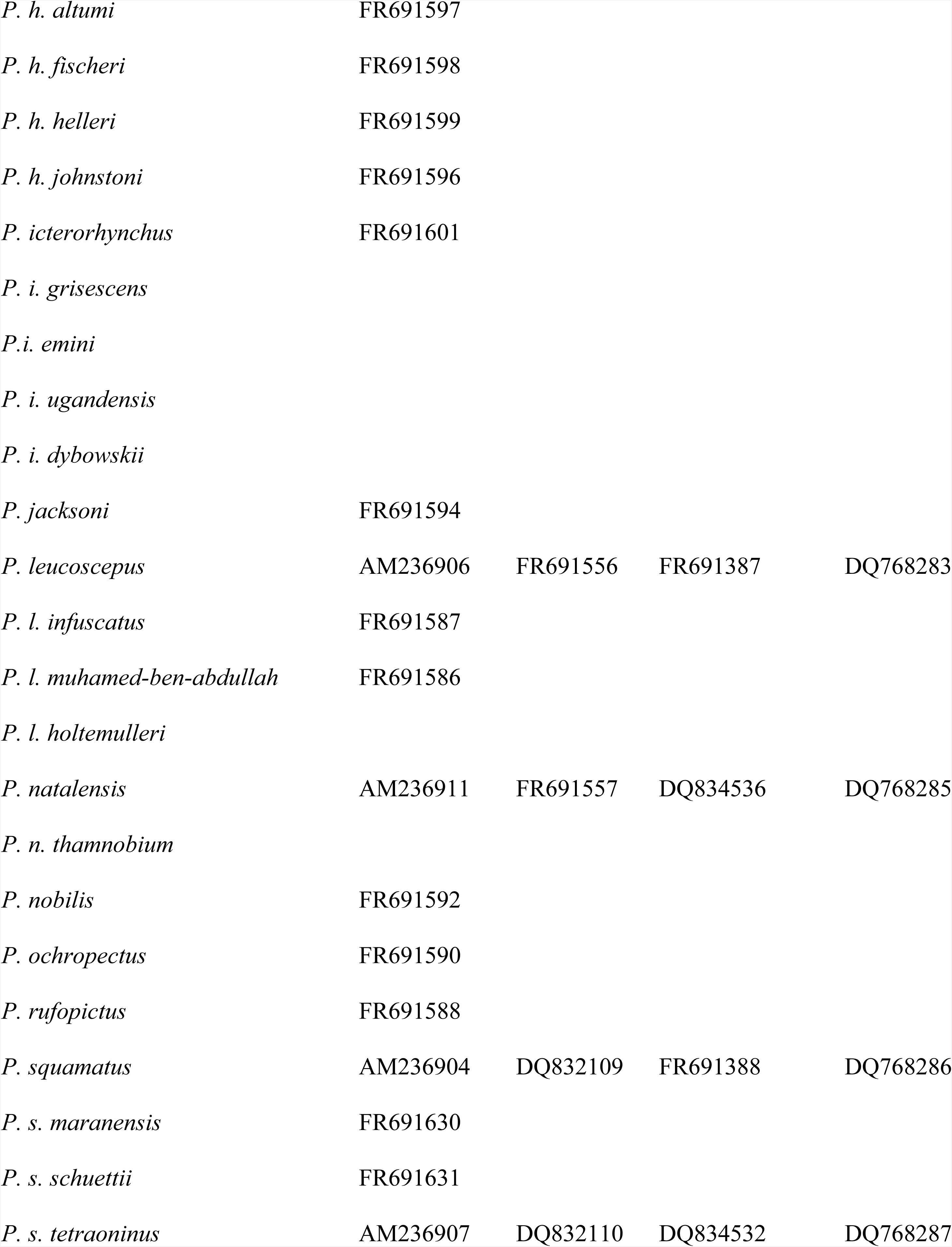
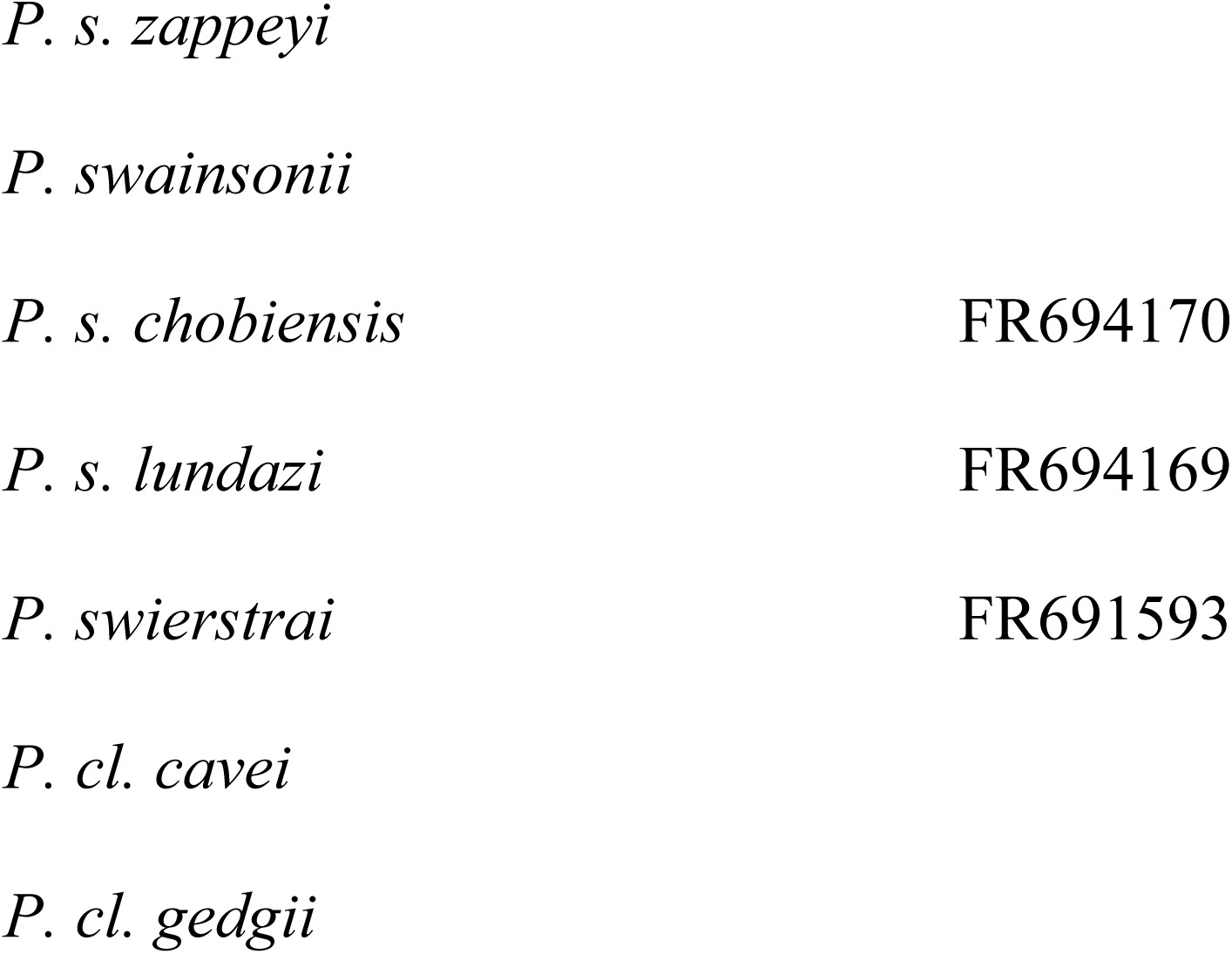
Spurfowl taxa examined and, where relevant, Genbank accession numbers for taxa sequenced for different molecular markers.

**Appendix 2.** Revised classification and common names for spurfowls based on multiple lines of evidence presented in this study. Family: Phasianidae; sub-family: Coturnicinae

| Genus, species, subspecies | Common names |
| --- | --- |
| *Genus: Pternistis* |  |
| *P. hartlaubi* | Hartlaub's Spurfowl |
| *P. camerunensis* | Mount Cameroon Spurfowl |
| *P. nobilis* | Handsome Spurfowl |
| *P. swierstrai* | Swierstra's Spurfowl |
| *P. erckelii* | Djibouti Spurfowl |
| *P. ochropectus* | Djibouti Spurfowl |
| *P. castaneicollis castaneicollis* | Chestnut-naped Spurfowl |
| *P. c. atrifrons* | Black-fronted Spurfowl |
| *P. jacksoni* | Jackson's Spurfowl |
| *P. squamatus* | Scaly Spurfowl |
| *P. schuetti schuettii* | Schuett's Spurfowl |
| *P. s. maranensis* |  |
| *P. ahantensis* | Ahanta Spurfowl |
| *P. griseostriatus* | Grey-striped Spurfowl |
| *P. hildebrandti hildebrandti* | Hildebrandt's Spurfowl |
| *P. h. fischeri* |  |
| *P. natalensis* | Natal Spurfowl |
| *P. adspersus* | Red-billed Spurfowl |
| *P. capensis* | Cape Spurfowl |
| *P. icterorhynchus* | Heuglin's Spurfowl |
| *P. bicalcaratus bicalcaratus* | Double-spurred Spurfowl |
| *P. b. ayesha* | Moroccan Spurfowl |
| *P. b. adamauae* |  |
| *P. harwoodi* | Harwood's Spurfowl |
| *P. clappertoni clappertoni* | Clapperton's Spurfowl |
| *P. c. sharpii* |  |
| *P. leucoscepus leucoscepus* | Yellow-necked Spurfowl |
| *P. l. infuscatus* |  |
| *P. swainsonii* | Swainson's Spurfowl |
| *P. cranchii* * | Cranch's Spurfowl |
| *P. rufopictus* | Grey-breasted Spurfowl |
| *P. afer afer* | Red-necked Spurfowl |
| *P. a. castaneiventer* |  |
| *P. a. humboldtii* |  |

\* subspecies elevated to species

## References

1. Gill, F and Donsker D (Eds). IOC World Bird List. doi:10.14344/IOC.ML.7.2 - http://www.worldbirdnames.org/-internetonly

2. Hall BP. The francolins, a study in speciation. Bull. Br. Mus. Nat. Hist. Zool. (Natural History). 1963. 10: 105–204.

3. Kimball RT, Braun EL, Zwartjes PW, Crowe TM., and Ligon JD. A molecular phylogeny of the pheasants and partridges suggests that these lineages are not monophyletic. Mol. Phylogenet. Evol. 1999; 11(1): 38–54.

4. Bock WJ, and Farrand J. The number of species and genera of recent birds. American Museum Novitates. 1980; 2703, 29 pp.

5. Milstein P le S, and Wolff SW. The over-simplification of our “francolins”.S. Afr. J. Wild. Res. Supplement 1, 1987; 58–65.

6. Crowe TM, and Little RM. Francolins, partridges and spurfowls: what’s in a name? Ostrich. 2004; 75: 199–203.

7. Johnsgard PA. The Quails, Partridges, and Francolins of the World. Oxford University Press, Oxford. 1988.

8. Madge S, and McGowan P. Pheasants, Partridges, and Grouse: A Guide to the Pheasants, Partridges, Quails, Grouse, Guineafowl, Buttonquails, and Sandgrouse of the world. Princeton Univ. Press, Princeton, N.J. 2002.

9. Little R and Crowe T. Gamebirds of southern Africa. 2nd edition. Cape Town: Random House Struik. 2011. Pp 135.

10. Sibley CG, and Monroe Jr BL. Distribution and Taxonomy of Birds of the World. Yale University Press, New Haven. 1990.

11. del Hoyo J, Elliott A, and Sargatal J. Handbook of the Birds of the World: New world Vultures to Guineafowl. Lynx edicions, Barcelona. 1994; Vol. II.

12. Roberts A. Synoptic check list of the birds of South Africa. Ann. Transvaal Mus. Vol. X. South Africa. 1924.

13. Roberts A. The Birds of South Africa. 1st ed. Riverside Pres. Edinburgh. 1940.

14. Wolters HE. Die Vogelarten der Erde. Paul Parey, Hamburg and Berlin. 1972–82.

15. Mandiwana-Neudani TG, Kopuchian C, Louw G, and Crowe TM. A study of gross morphological and histological syringeal features of true francolins (Galliformes: *Francolinus, Scleroptila, Peliperdix* and *Dendroperdix* spp.) and spurfowls (*Pternistis* spp.) in a phylogenetic context. Ostrich. 2011; 82: 115–127.

16. van Niekerk JH, and Mandiwana-Neudani TG. The phylogeny of francolins (Francolinus Dendroperdix, Peliperdix and Scleroptila) and spurfowls (Pternistis) based on chick plumage (Galliformes: Phasianidae. Avian Research 2018; 9:2.

17. Mandiwana-Neudani TG, Bowie RCK, Hausberger M, Henry L, and Crowe TM. Taxonomic and phylogenetic utility of variation in advertising calls of francolins and spurfowls (Galliformes: Phasianidae). Afr. Zool. 2014; 49: 54–82.

18. van Niekerk JH. Vocal structure, behavior and partitioning of all 23 *Pternistis* spp. into homologous sound (and monophyletic) groups. Chinese Birds. 2013; 4(3): 210–231. DOI 10.5122/cbirds.2013.0020

19. Bloomer P, and Crowe TM. Francolin phylogenetics: molecular, morphobehavioral, and combined evidence. Molecular Phylogenetics and Evolution. 1998; 9(2): 236–254.

20. Crowe TM, Bowie RCK, Bloomer P, Mandiwana TG, Hedderson TAJ, Randi E, et al. Phylogenetics, biogeography and classification of, and character evolution in, gamebirds (Aves: Galliformes): Effects of character exclusion, data partitioning and missing data. Cladistics. 2006; 22: 1–38.

21. Chapin JP. A new genus *Acentrortyx* proposed for *Francolinus nahani* Dubois. The Auk. 1926; 43: 235.

22. Cohen C, Wakeling J L, Mandiwana-Neudani TG, Sande E, Dranzoa C, Crowe TM, et al. Phylogenetic affinities of evolutionarily enigmatic African galliforms: the Stone Partridge *Ptilopachus petrosus* and Nahan’s Francolin *Francolinus nahani*, and support for their sister relationship with New World quails. Ibis. 2012; 154: 768–780.

23. Bowie RCK, Cohen C, and Crowe TM. Ptilopachinae: a new subfamily of the Odontophoridae (Aves: Galliformes). Zootaxa. 2013; 3670: 097–098.

24. Zachos FE. Species Concepts in Biology. Historical Development, Theoretical Foundations and Practical Relevance. Springer International Publishing Switzerland, [place not stated]. 2016.

25. Winker K. Chapter 1: Subspecies represent geographically partitioned variation, a gold mine of evolutionary biology, and a challenge for conservation. Ornithol. Monogr. 2010; 67: 6–23.

26. De Queirroz K. Species concepts and species delimitation. Syst Biol. 2007; 56(6): 879–886.

27. Linder HP, de Klerk HM, Born J, Burgess ND, Fjeldså J, and Rahbek C. The partitioning of Africa: statistically defined biogeographical regions in sub-Saharan Africa. J. Biogeogr. 2014; 39: 1189–1205.

28. Pruett CL, and Winker T. Alaska song sparrows (*Melospiza melodia*) demonstrate that genetic marker and method of analysis matter in subspecies assessments. The American Ornithologists’ Union. 2010; 67: 162–171.

29. McCarthy EM. Handbook of avian hybrids of the world. Oxford University Press, New York. 2006.

30. Casacci LP, Barbero F, and Balletto E. The “Evolutionarily Significant Unit” concept and its applicability in biological conservation. Italian Journal of Zoology. 2014; 81 (2): 182–193. https://doi.org/10.1080/11250003.2013.870240

31. Crowe TM, Essop MF, Allan DG, Brooke RK, and Komen J. Overlooked units of comparative and conservation biology: the Black Korhaan *Eupodotis afra* (Otididae) as a case study. Ibis. 1994; 136: 166–175.

32. Mallet J. Hybridization as an invasion of the genome. Trends Ecol. Evol. 2005; 20: 229–237.

33. Mallet J. Hybridization, ecological races and the nature of species: empirical evidence for the ease of speciation. Philosophical Transactions of the Royal Society B: Biological Sciences. 2008; 363: 2971–2986.

34. Crowe TM. The evolution of guinea-fowl (Galliformes, Phasianidae, Numidinae) taxonomy, phylogeny, speciation and biogeography. Annals of the South African Museum. 1978; 76: 43– 136.

35. Bowie RCK, Fjeldså J, Kiure J, and Kristensen J. A new member of the greater double-collared sunbird complex (Passeriformes: Nectariniidae) from the Eastern Arc Mountains of Africa. Zootaxa. 2016; 4175: 23–43.

36. Crowe TM, Harley EH, Jakutowicz MB, Komen J, and Crowe AA. Phylogenetic, taxonomic and biogeographical implications of genetic, morphological, and behavioural variation in francolins (Phasianidae. *Francolinus*). Auk. 1992; 109(1): 24–42.

37. Edwards SV, and Wilson AC. Phylogenetically informative length polymorphism and sequence variability in mitochondrial DNA of Australian songbirds (*Pomatostomus*). Genetics. 1990; 126: 695–711.

38. Fumihito A, Miyake T, Takada M, Ohno S, and Kondo N. The genetic link between the Chinese bamboo partridge (*Bambusicola thoracica*) and the chicken and junglefowls of the genus *Gallus*. Proceedings of the National Academy of Sciences of the United States of America. 1995; 92: 11053–11056.

39. Moum T, Johansen S, Erikstad KE, and Piatt JF. Phylogeny and evolution of the auks (subfamily Alcinae) based on mitochondrial DNA sequences. Proceedings of the National Academy of Sciences of the United States of America. 1994; 91: 7912–7916.

40. Sorenson MD, Ast JC, Dimcheff DE, Yuri T, and Mindell DP. Primers for a PCR-based approach to mitochondrial genome sequencing in birds and other vertebrates. Mol. Phylogenet. Evol. 1999; 12: 105–114.

41. Armstrong MH, Braun EL, and Kimball RT. Phylogenetic utility of avian ovomucoid intron G: a comparison of nuclear and mitochondrial phylogenetics in Galliformes. Auk. 2001; 118: 799– 804.

42. Friesen VL, Congdon BC, Walsh HE, and Birt TP. Intron variation in Marbled Murrelets detected using analyses of single-stranded conformation polymorphisms. Mol. Ecol. 1997; 6: 1047–1058.

43. Primmer CR, Borge T, Lindell J, and Saetre GP. Single nucleotide polymorphism characterization in species with limited available sequence information: high nucleotide diversity revealed in the avian genome. Mol. Ecol. 2002; 11: 603–612.

44. Kornegay JR, Kocher TD, Williams LA, and Wilson AC. Pathways of lysozyme evolution inferred from the sequences of cytochrome b in birds. J. Mol. Evol. 1993; 37: 367–379.

45. Edwards SV, Arctander P, and Wilson AC. Mitochondrial resolution of a deep branch in the genealogical tree for perching birds. Proceedings of the Royal Society. 1991; 243: 99–107.

46. Hennig W. Phylogenetic Systematics, translated by D. Davis and R. Zangerl, Urbana: University of Illinois Press, 1966 (reprinted 1979).

47. Kluge AG. Total evidence or taxonomic congruence: cladistics or consensus classification. Cladistics. 1998; 14: 151–158.

48. Nixon KC, and Carpenter J M. On simultaneous analysis. Cladistics. 1996, 12: 221–241.

49. Farris JS. The logical basis of phylogenetic analysis. In Platnick, Norman I.; Funk, Vicki. Advances in cladistics, vol. 2, Columbia University Press, New York, 1983; pp 7–36.

50. Rindal E, Brower AVZ. Do model-based phylogenetic analyses perform better than parsimony? A test with empirical data. Cladistics. 2011; 27: 1–4.

51. Snow DW. An Atlas of Speciation in African Non-passerine birds. London: British Museum. 1978.

52. Harrison JA, Allan DG, Underhill LG, Herremans M, Tree A J, Parker V, et al. The Atlas of Southern African Birds: Non-passerines. Vol. 1. Johannesburg: BirdLife South Africa. 1997.

53. Komen J. Preliminary observations of the social pattern, behaviour and vocalisations of Hartlaub’s Francolin. S. Afr. J. Wildl. Res. 1987; 1: 82–86.

54. Little R. Terrestrial Gamebirds and Snipes of Africa. Jacana Media, Johannesburg, South Africa. 2016a. pp 301.

55. Töpfer T, Podsiadlowski L, and Gedeon K. Rediscovery of the Black-fronted Francolin *Pternistis (castaneicollis) atrifrons* (Conover, 1930) (Aves: Galliformes: Phasianidae) with notes on biology, taxonomy and conservation. Vertebrate Zoology. 2014; 64: 261–271.

56. Moreau, RE. The bird faunas of Africa and its islands. New. York and London, Academic Press. 424 pages. 1966.

57. Balinsky BI. Patterns of animal distribution on the: African continent. Ann. Cape Prov. Mus. 1962; 2: 299–310.

58. Bobe R. The evolution of arid ecosystems in eastern Africa. J. Arid Environ. 2006, 66: 564–584.

59. Vernon CJ. Biogeography, endemism and diversity of animals in the Karoo. In Dean, W.R.J.and Milton, S.J. editors. The Karoo. Ecological patterns and processes. Cambridge University Press, Cambridge. 1999; 57–78.

60. Irwin MPS, Benson CW, and White CMN. The significance of valleys as avian zoogeographical barriers. Ann. Cape Prov. Mus. 1962; 2: 155–189.

61. Crowe TM, and Crowe A A. The genus *Francolinus* as a model for avian evolution and biogeography in Africa. Pp. 207–231. In: Proceedings of the International Symposium on African vertebrates (Schuchmann KL, ed.). Museum Alexander Koenig, Bonn. 1985.

62. Crowe TM and Crowe AA. Patterns of distribution, diversity and endemism in Afrotropical birds. London. J. Zool. 1982; 198: 417–442.

63. Cotterill FPD. Geomorphological influences on vicariant evolution in some African mammals in the Zambezi basin: some lessons for conservation. In: Ecology and Conservation of Small Antelope. Proceedings of an International Symposium on Duiker and Dwarf Antelope in Africa, Chapter: Geomorphological influences on vicariant evolution in some African mammals in the Zambezi basin: some lessons for conservation, Publisher: Filander Verlag, Fürth, Editors: A. Plowman. 2003; 11–58.

64. Roberts AA. New species of *Pternistis*from Salisbury, Rhodesia. Ostrich. 1947; 18(2): 197.

65. Little R. Hybrid spurfowl unravelled. African Birdlife Letters November/December. 2016b; 5(1): 6–8.

66. Little R. Mix & match Hybrid spurfowl. African Birdlife News & Views July/August. 2016c; 4(5): 16.

